# Multiple-omics analyses of the *Alviniconcha* holobiont reveal multi-faceted adaptations to deep-sea hydrothermal vents

**DOI:** 10.1101/2025.05.21.655246

**Authors:** Hui Wang, Yuran Dai, Chong Chen, Xing He, Menggong Li, Yadong Zhou, Jack Chi-Ho Ip, Jin Sun

**Author notes:** Corresponding Author: Jin Sun.

## Abstract

Deep-sea hydrothermal vents are ‘extreme’ environments with constantly fluctuating physicochemical conditions – but dense animal aggregations thrive primarily through symbiosis with chemoautotrophic bacteria to exploit the unusual chemistry. *Alviniconcha* snails, which harbor symbionts in their enlarged gill at an intracellular-extracellular intermediate state, are a prime example. Here, we present chromosome-level genomes of two *Alviniconcha* species (*A. adamantis* and *A. marisindica*) to investigate the adaptations of this holobiont. Significant expansion of solute carrier families enhances nutrient transport between the two parties. *Alviniconcha* lacks complete methionine biosynthesis pathways, compensated by symbiont provisioning, highlighting host-symbiont metabolic complementarity. High myoglobin expression in gills, contradicting prior reports of hemoglobin, suggests myoglobin-mediated oxygen storage to mitigate fluctuating environmental oxygen. Spatial transcriptomics further delineated gill’s functional zones on the gill filament responsible for symbiont digestion via phagocytosis in bacteriocytes, oxygen transport in secretory zones, and ciliary water flow regulation. Our findings elucidate molecular and physiological adaptations underpinning the *Alviniconcha* holobiont’s success in dynamic and treacherous vent ecosystems.

## INTRODUCTION

Symbiosis is a highly successful adaptive strategy for animals to thrive in extreme environments (Apprill, 2020). Chemosymbiosis refers to the mutually beneficial relationship between host animals and chemoautotrophic bacteria, which evolved independently many times in environments rich in hydrogen sulfide and other reduced compounds, such as hydrothermal vents, hydrocarbon seeps, and organic falls (Dubilier et al., 2008; Sogin et al., 2020). These host animals harbor specific lineages of bacteria in specialized organs such as gill, trophosome, or esophageal gland where the bacteria can safely multiply; while they provide energy and nutrition to the host by the oxidation of reduced substrates, such as hydrogen sulfide, hydrogen, and methane, to support the host and its growth (Sogin et al., 2021). The ambient environment around hydrothermal vents typically exhibit low oxygen concentrations and high concentrations of reducing substances, supporting large animal communities (Dubilier et al., 2008), where the most dominant megafaunal animals are typically associated with symbionts for sustaining their biomass and survival (Dover, 2000). Annelid tubeworms in the family Siboglinidae (Dubilier et al., 2008) and several groups of molluscs, such as gastropods in Abyssochrysoidea (Beinart et al., 2012; Suzuki et al., 2005) and Peltospiridae (Goffredi et al., 2004; Nakagawa et al., 2014), as well as bivalves in Vesicomyidae and Bathymodiolinae are groups that include chemosymbiotic animals commonly found in vents. The host and its symbionts, collectively known as the ‘holobiont’, have co-developed multifaceted ways to cope with each other as a key adaptation to exploit the chemosynthetic potential of the vent environment.

Gastropod molluscs are one of the most abundant and diverse animal groups in deep-sea chemosynthetic communities. *Alviniconcha* (Abyssochrysoidea: Paskentanidae) is a vent-endemic genus characterized by a hairy periostracum, widely distributed in Pacific and Indian oceans. This genus includes six species (Johnson et al., 2014): *A. adamantis*, *A. marisindica*, *A. kojimai*, *A. hessleri*, *A. boucheti*, and *A. strummeri*. Their intestines are dramatically reduced; these snails host chemoautotrophic symbionts in their enlarged gill, which is the main source of nutrition for the snails. The symbionts are housed in bacteriocytes occupying a specific region of the gill filaments called the bacteriocyte zone (Laming et al., 2020).

Compared to other chemosymbiotic molluscs, such as vesicomyid clams and peltospirid snails which have true endosymbionts fully contained within animal cells, the symbiosis mode in *Alviniconcha* seems to be more versatile. First of all, the symbionts are at the intermediate phase between the intracellular and extracellular stages (Endow and Ohta, 1989). The symbionts are contained in narrow tube-like indentations of the bacteriocyte surface which open apically to the external environment, akin to the situation in bathymodioline mussels (Ikuta et al., 2021). Secondly, *Alviniconcha* snails acquire symbionts from the ambient environment by horizontal transmission (Breusing et al., 2022a), and different species prefer different symbiont lineages. For example, *A. adamantis* harbors sulfur-oxidizing Gammaproteobacteria, while *A. marisindica* harbors Campylobacterota with the capacity to use hydrogen sulfide and hydrogen gas (Breusing et al., 2022a; Miyazaki et al., 2020). Different symbionts of *Alviniconcha* display strain-level flexibility influenced by environmental factors, enabling them to adapt to the dynamic conditions of their ‘extreme’ vent habitats, and it is a good example of symbiont-driven ecological niche partitioning (Breusing et al., 2022a). *Alviniconcha* species are typically found right next to the active vent edifice, implying that they are heavily influenced by the vent chemistry. However, we still lack a holistic understanding on how the host and symbiont in the *Alviniconcha* holobiont work together to conquer and thrive in the ‘extreme’ hydrothermal vent environment.

Here, we combine multiple-omics methods to study the *Alviniconcha* holobiont system. In order to unravel the interactions between host and symbionts, we investigate the genomic features from both host and symbiont genomes, and employ cutting-edge spatial transcriptomic analysis on their gene expression patterns. We aim to gain insights into how the host cooperates with symbionts to maintain mutualism and the molecular adaptations in the whole holobiont that underpin its success.

## RESULTS AND DISCUSSION

### Genomic features of the two *Alviniconcha* species

We assembled high-quality genomic contigs of *Alviniconcha marisindica* and *A. adamantis* with the combination of Oxford Nanopore Technologies long-read and Illumina sequencing techniques. The statistics of the newly assembled genomes are summarized in Table S1. These contig-level genomes were scaffolded into chromosome-level genome assemblies using Hi-C sequencing, uncovering 18 chromosomes in both species. The final chromosome-level genome size was 825 Mb for *A. marisindica* and 795 Mb for *A. adamantis* (Figure S1). Combining *ab initio*, transcripts, and homology evidence, a total of 25,691 and 23,163 predicted protein-coding genes were annotated in *A. marisindica* and *A. adamantis*, respectively; with BUSCO scores of 95.7% and 96.1%, respectively (Table S1 in Supplementary Table).

For repetitive analysis, we identified a large amount of interspersed repeat content in their genomes (32.1% for *A. marisindica* and 34.9% for *A. adamantis*). Moreover, both genomes contain a substantial number of unclassified repeats (22.5% in *A. marisindica* and 18.3% in *A. adamantis*) and tandem repeats (31.6% in *A. marisindica* and 12.0% in *A. adamantis*) (Table S2 in Supplementary Table). Comparing genomic synteny in *Alviniconcha* to the Molluscan Linkage Groups (MLGs) (Sigwart et al., 2025) revealed two independent fusion events, including Chr01 is formed by the fusion of MLG01 and MLG19 and Chr02 originating from the fusion of MLG10 and MLG14, suggesting a high karyotype conservation within genus *Alviniconcha* (Figure 1B). To determine the phylogenetic positions of the *Alviniconcha* in Gastropoda, all trees generated using IQ-TREE MFP model, LG+C60 model and phylobayes with different matrices congruently placed the two *Alviniconcha* species in the subclass Caenogastropoda with maximum support, among other taxa currently assigned to the order Littorinimorpha (Figure S2 and Table S3 in Supplementary Table). Like other recent phylogenetic studies of Caenogastropoda, our tree also recovered a paraphyletic Littorinimorpha (Wang et al., 2024b).

**Figure 1.**
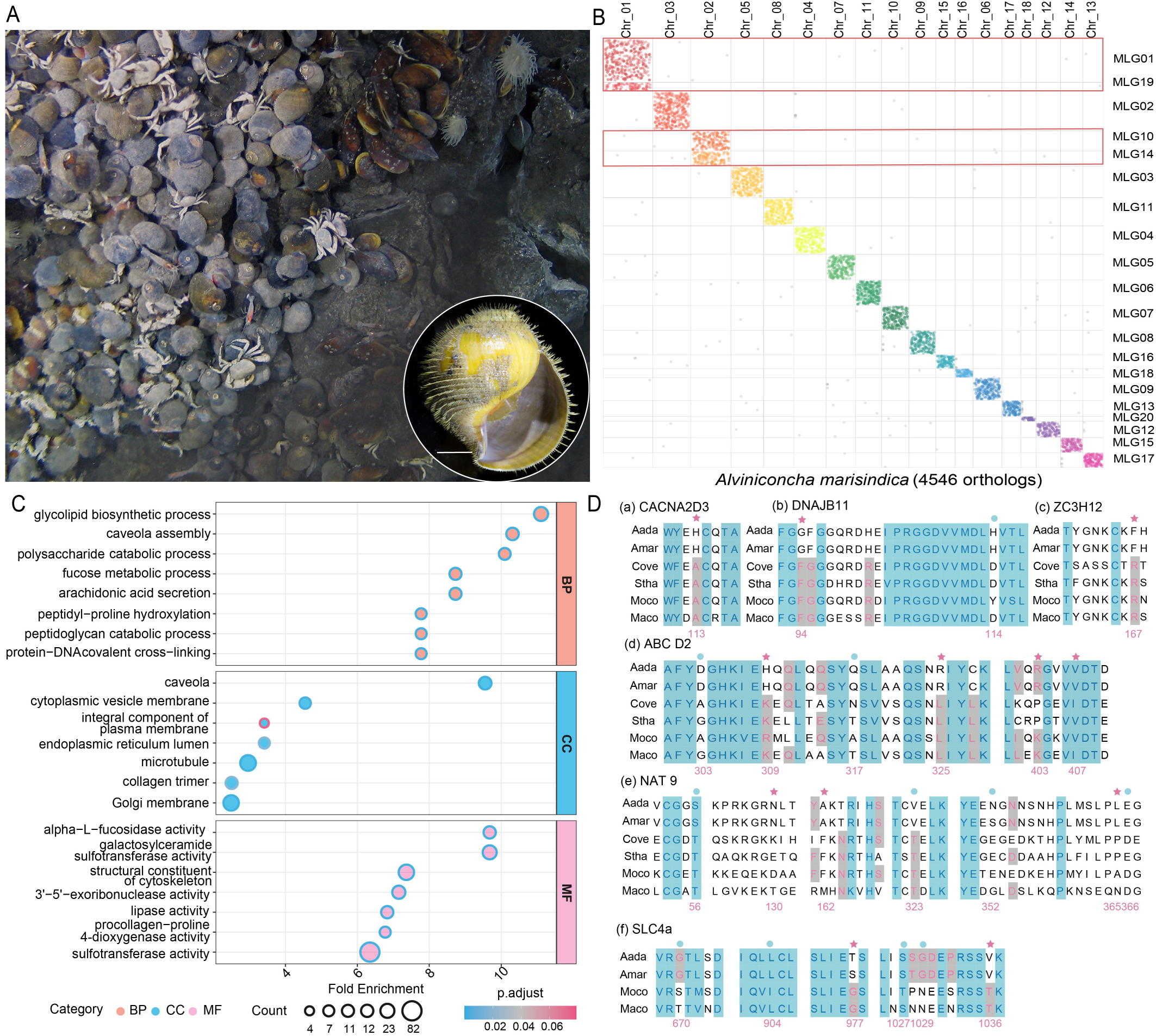
The genomic features of *Alviniconcha marisindica* with comparative syntenic analysis, gene family analysis and positively selected genes. A, *A. marisindica* occurs in high abundance in the Wocan vent field with a specimen photograph in the lower-right corner. Scale bar: 1cm. B, Oxford plots comparing gene occupancy of *A. marisindica* (horizontal) to the ancestral molluscan linkage groups (MLGs, vertical). C, GO enrichment analysis of the expanded gene families GO terms in *Alviniconcha*, those exhibiting statistically significant differences are shown. D, Sequence alignment of positively selected genes. Pink asterisk indicates that the amino acids in *Alviniconcha* spp. have a posterior probability higher than 95%, and blue dot indicates that the sites have a posterior probability between 90-95%. Abbreviasions: Aada: *A. adamantis*; Amar: *A. marisindica*; Moco: *Monoplex corrugatus*; and Maco: *Marisa cornuarietis; Stha: Stramonita haemastoma; Cove: Conus ventricosus*.

### Gene family analyses and positive selection genes

We detected the significant expansion of 213 gene families and contraction of 304 gene families in the two *Alviniconcha* species (Table S4 in Supplementary Table and Figure S3). These expanded gene families are involved in transmembrane transporter activity, sulfotransferase activity, and hydrolase activity etc., indicative of their contributions to nutrient transport and metabolism of these two species to cope with the symbionts (FDR <0.05, Figure 1C). Moreover, the expansion of solute carriers (SLC), including SLC5, SLC13, SLC21 and SLC46, in *Alviniconcha* may play a critical role in host-symbiont interaction by regulating the transportation of diverse substances between hosts and symbionts (Höglund et al., 2011; Pizzagalli et al., 2021; Verri et al., 2012), such as sodium/hydrogen ions, organic anions, glucose, monocarboxylate, steroid sulfate, and large neutral amino acids.

Gene ontology (GO) enrichment analysis of the contracted gene families revealed a reduction in multiple cellular activities (Figure S4). Among them, methionine biosynthesis is a conserved pathway in molluscs, but this pathway is significantly contracted in *Alviniconcha* with the largest decreased fold changes compared to other molluscs, which may be related to the fact that the symbionts provide methionine to the host as suggested by the genome of the symbionts (Breusing et al., 2022b; Yang et al., 2020). The lack of an obligate methionine biosynthesis enzyme (cysteine-S-conjugate beta-lyase, metC) in both *Alviniconcha* species suggests an incomplete methionine biosynthesis pathway, and a likely dependency of this obligate amino acid on the symbiont. A similar finding was previously reported from the coral *Acropora digitifera* whose genome lacks a key enzyme for cysteine synthesis (Shinzato et al., 2011). The energetic and nutritional requirements of the *Alviniconcha* snails are heavily reliant on the symbionts, as the intestine is largely reduced. In addition, the G protein-coupled receptor signaling pathway, cell surface receptor signaling pathway, and Wnt signaling pathway related to controlling cell-to-cell interactions, were also contracted and enriched in *Alviniconcha*. The reductions of these gene families are linked with the sensing of chemical cues, shown to be crucial to shallow-water and freshwater gastropods (Fu et al., 2022; Rondón et al., 2024; Sun et al., 2019). Altogether, these suggest *Alviniconcha* snails have reduced capacity in detecting various chemical cues commonly present in non-chemosynthetic shallow-water and freshwater environments.

To identify genes under positive selection in *Alviniconcha* species, three models (two branch-site model and one site model) were used to compare them with shallow-water marine gastropods or freshwater snails, finding 78 genes positively selected in two *Alviniconcha* (*P* <0.05). These genes are likely related to *Alviniconcha*’s adaptation to the vent environment (Table S5 in Supplementary Table), which includes sodium-driven chloride bicarbonate exchanger 4 (SLC4) participating in ions exchange, voltage-dependent calcium channel, ATP-binding cassette sub-family D involved in ATP binding, N-acetyltransferase 9 contributing to acyltransferase activity, and dnaJ homolog subfamily B member 1 serving as molecular chaperones (Figure 1D).

### Myoglobin, not hemoglobin

While *Alviniconcha* uses hemocyanin as the primary oxygen carrier, it was reported to exhibit a high concentration of hemoglobin specifically in the gill tissue, hypothesized to function as additional oxygen carriers to bacteriocytes, as known from vesicomyid clam and tubeworm symbionts *(Wittenberg and Stein, 1995)*. Analyzing the genome of two *Alviniconcha* species, we failed to recover any homologue of hemoglobin, while two copies of proteins annotated as myoglobin were detected at very high protein expression levels in our gill proteomic analysis (intensity: 4.54x10^11^, among the top 30 most abundant proteins in gill) and gill gene expression (table S6 in Supplementary Table). To confirm the identity of these proteins as myoglobins, a phylogenetic analysis was performed on the known globins as well as hemoglobins, showing these *Alviniconcha* myoglobins were clustered with other myoglobins, such as those from the gastropods *Cerithidea rhizophorarum* (P02215) and *Buccinum undatum* (Q7M424) (Figure 2A, Figure S5A). To further examine their structure similarity, a 3D structure comparison was performed by AlphaFold3 and DALI structure comparison servers. These results show that the *Alvinconcha* myoglobins are closer to other known myoglobins rather than hemoglobins (Figure 2B). The gene localization of these myoglobins in *Alviniconcha* gills confirms the local expression of these myoglobins in the gill filaments (Figure 2C and Figure S5B). Our results therefore suggest that the previous detection of ‘hemoglobin’ in the gill of *Alviniconcha* was a misidentification of myoglobins, which are present in high abundance.

**Figure 2.**
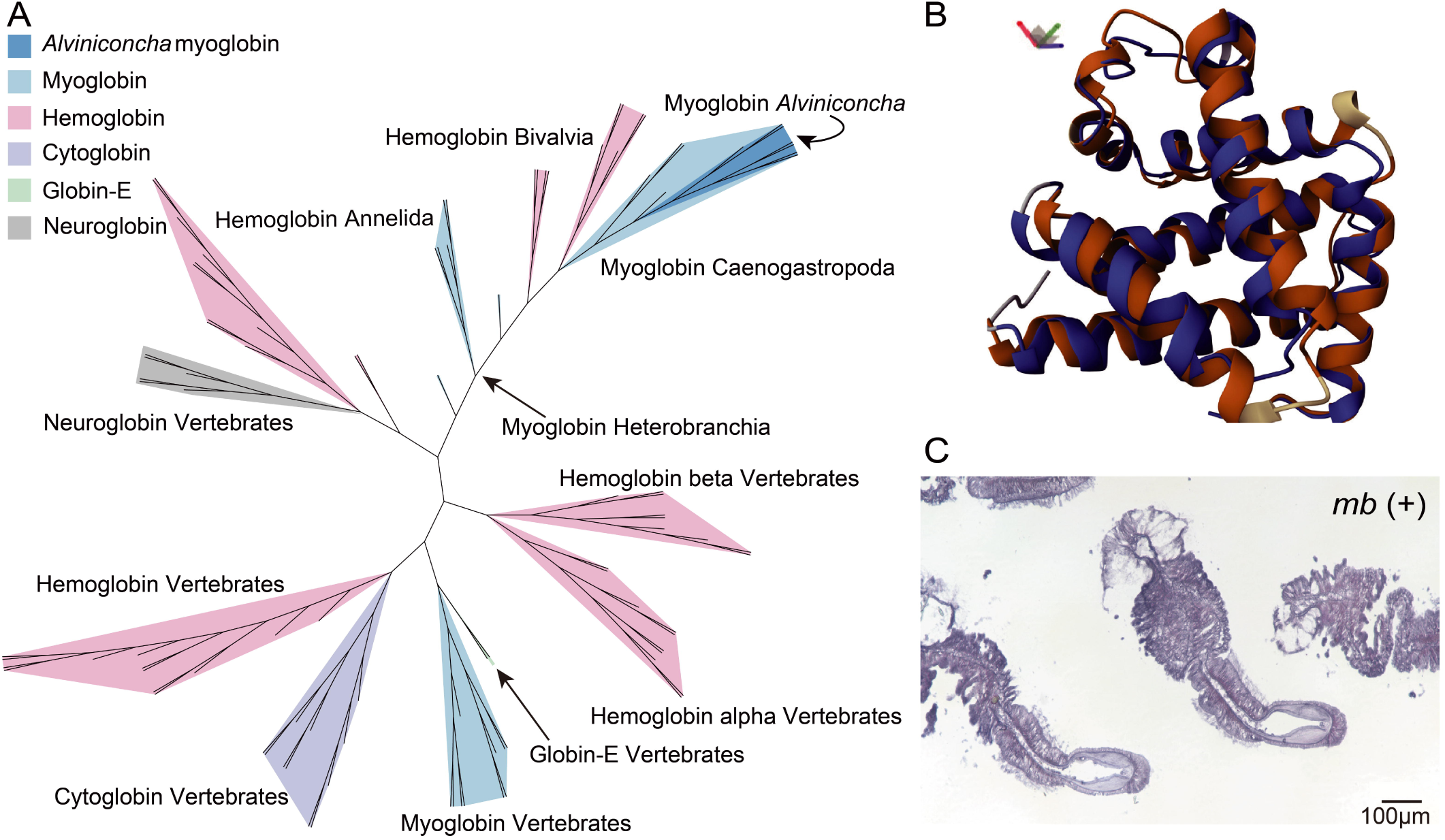
Identified myoglobin in *Alviniconcha*. A, The unrooted tree was constructed with myoglobin in *Alviniconcha* and other typical globin proteins. B, Protein structure as predicted by AlphaFold and shown using the RCSB Protein Data Bank. Compared with whale (*Balaenoptera borealis*) myoglobin for 3D structures comparison. C, The ISH of myoglobin in *A. marisindica* gill section. Scale bar: 100 µm.

*Alviniconcha* snails live near vent orifices where many biochemical factors (such as dissolved oxygen levels and H_2_S concentrations) change rapidly due to the mixing between hot anoxic vent fluid with cold oxygen-rich bottom water (Henry et al., 2008; Laming et al., 2020; Trembath-Reichert et al., 2019). The symbionts rely on the oxidation of H_2_S for carbon fixation through chemosynthesis, while the host itself also has a high demand for oxygen for survival. High concentrations of HLS and oxygen exhibit an inverse relationship, with their spatial-temporal availability fluctuating dynamically (Watanabe and Kojima, 2015). Compared to hemoglobins, myoglobins display higher binding affinities for oxygen (Aharoni and Tobi, 2019). The myoglobins likely serve as the oxygen storage. *Alviniconcha* snails can thus extract and store oxygen while they are bathed by relatively oxygen-rich waters (i.e., mixing of more bottom water than vent fluid), and use these for respiration and chemosynthesis when surrounded by anoxic conditions (i.e., mixing of more vent fluid and less bottom water). High concertation of myoglobin is an adaptation strategy in marine mammals, such as whales and seals, for their deep-sea diving (Wright and Davis, 2015), in comparison with terrestrial mammals.

### Symbiont localization and genome features

To confirm the location of the symbionts in *Alviniconcha* gill tissues, fluorescent *in situ* hybridization (FISH) was carried out on transverse sections of the gill filament of *A. marisindica*, yielding positive signals of bacteria only in the bacteriocytes zone (BZ), which consistent with previous studies (Laming et al., 2020; Miyazaki et al., 2020) (Fig 3A). FISH also identified the ciliated zone (CZ) with characteristic of extensive cilia and secretory zone (SZ) (details in Spatial Transcriptome section), whereas the supporting axis (SA) were only labelled by cell membrane specific ConA signal and DAPI signal.

**Figure 3.**
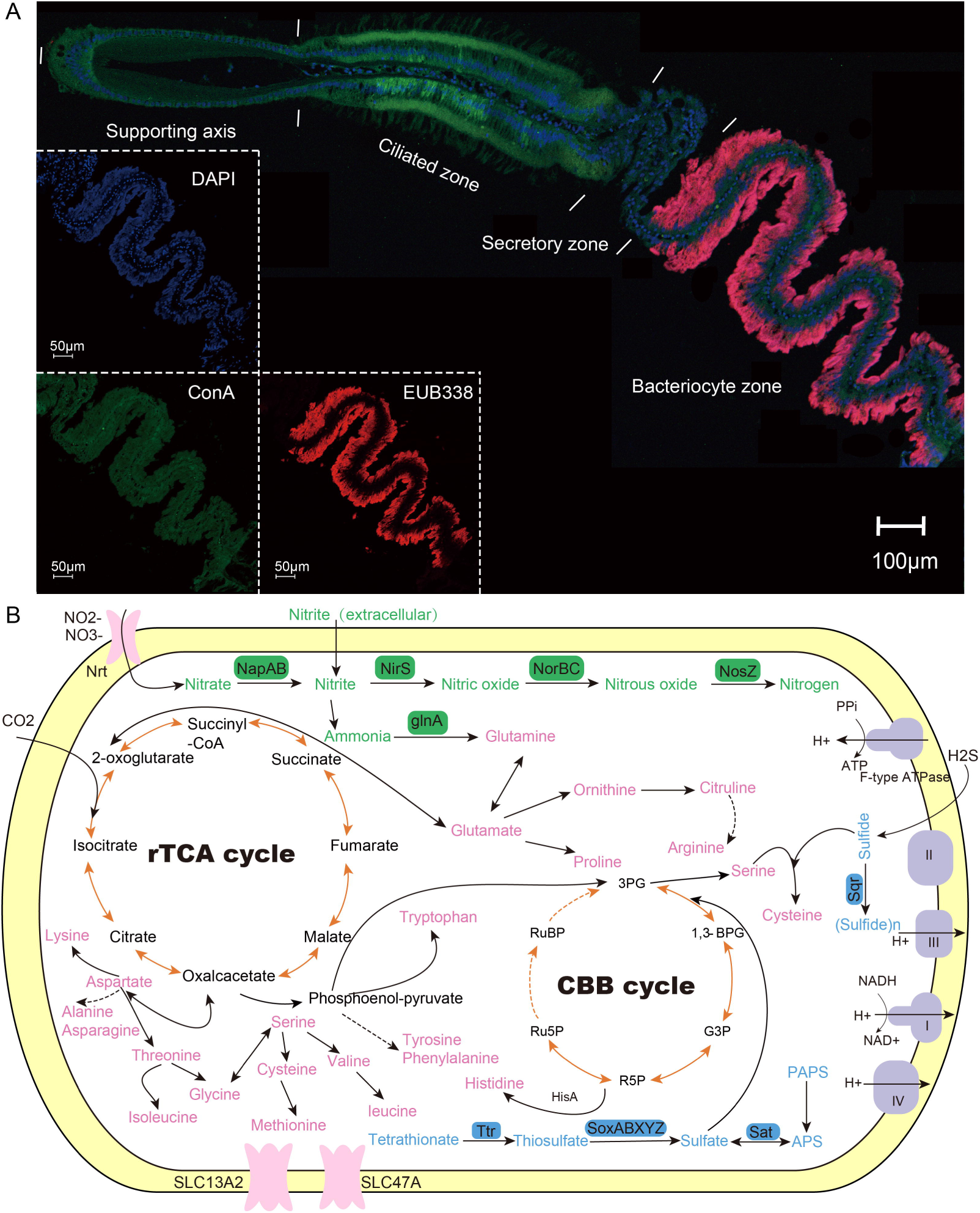
Symbionts in the gill of *Alviniconcha marisindica* and central metabolism of the symbionts of *A. marisindica.* A, FISH of *A. marisindica* with Cy5-labeled bacteria-specific probe EUB338, ConA staining cell membrane, DAPI staining nucleic acid. B, Overview of metabolic pathways in the *Candidatus* Sulfurovum *Alviniconcha* (*C*Sal). Solid arrows represent the genes or enzymes in the symbiont, whereas the dashed arrows indicate absence. Amino acids shown in solid arrows have a complete biosynthesis pathway, and incomplete pathways are shown in dashed arrows.

The genome of *Candidatus* Sulfurovum *Alviniconcha* (*C*Sal) was initially reported by Yang et al. as a preprint (Yang et al., 2020) but this remains unpublished in the peer-reviewed literature, and it is published for the first time here. ONT long read assembly recovered the *C*Sal genome has a length of 1.47 Mb (in two scaffolds, 98.16% completeness, 0.82% contamination) with 1429 predicted genes, 92.65% of which were successfully annotated. It possesses fewer coding sequences than other available sequenced genomes within the phylum, possibly because of genome streamlining (Table S7 in Supplementary Table). Flagellar or chemotaxis genes and many cell envelope biogenesis and non-essential metabolic genes are missing in the genome, suggesting a lack of a free-living lifestyle, like capsular polysaccharide biogenesis.

We generated new metaproteome data to study protein expression in the gill of *A. marisindica*, to further illustrate the host-symbiont interaction. The proteins extracted from the gill of three *A. marisindica* individuals were identified and quantified by liquid chromatography with tandem mass spectrometry (LC-MS/MS), resulting in 3087 host proteins and 207 symbiont proteins (for an overview of the total protein identifications in all samples see Table S6, Table S8 in Supplementary Table and Figure S6). Analysis of the core metabolic proteins of the symbiont genome reveals its versatile chemolithoautotrophic capacity. Firstly, *C*Sal contains the complete reductive tricarboxylic acid (rTCA) cycle for carbon fixation, with three proteins (KorA, KorB, and ICDH) highly expressed in the proteome. Secondly, this genome lacks the complete sulphide oxidation pathway, as indicated by the absence of genes in the DsrAB complex. The Sox multi-enzyme system (SoxX-SoxY-SoxZ-SoxA-SoxB) allows the generation of energy from thiosulphate oxidation. The absence of a sulphate/thiosulphate transporter in the *C*Sal genome suggests it can only use endogenous thiosulphate. *C*Sal can actively consume environmental sulphide, which is supported by highly expressed SQR (top 50 of the proteome) and *cysK* in the genome that are involved in the conversion of sulphide to polysulphides. In addition, *C*Sal lacks the sulphur globule protein genes (*sgp*) for intracellular sulphur storage, indicating that this symbiont is likely dependent on intracellular polysulphides for sulphur storage. Lastly, the nutrient synthesis pathways (such as amino acids and vitamins/cofactors) were diverse, with a total of 17 amino acids and 8 vitamins/cofactors that can be synthesized by the symbiont.

### Spatial transcriptomics shed light on functional partitioning of the gill filament

To reveal the spatial distribution of gene expressions and interactions of both the host and *C*Sal, we applied spatial transcriptomics on four separately dissected PFA-fixed gill tissues. Under bin 50 (50 x 50 DNBs, i.e. center-to-center distance of 25µm), the mean captured MID and genes per bin in the section were 223 and 191, respectively. The adjacent slide was HE stained, and the four areas in the gill (i.e. SA, CZ, SZ, and BZ) were distinguished (Figure 4A). The spatial distribution pattern of 16rRNA of *C*Sal was specifically detected in BZ, which was highly consistent with the former FISH result (Figure 3A), verifying the reliability of the data. Among these four gill tissues, gill No. IV exhibited higher symbiont density as characterized by the higher read counts mapped to the 16S rRNA, suggesting symbiont density heterogeneity among individuals (Figure 4B).

**Figure 4.**
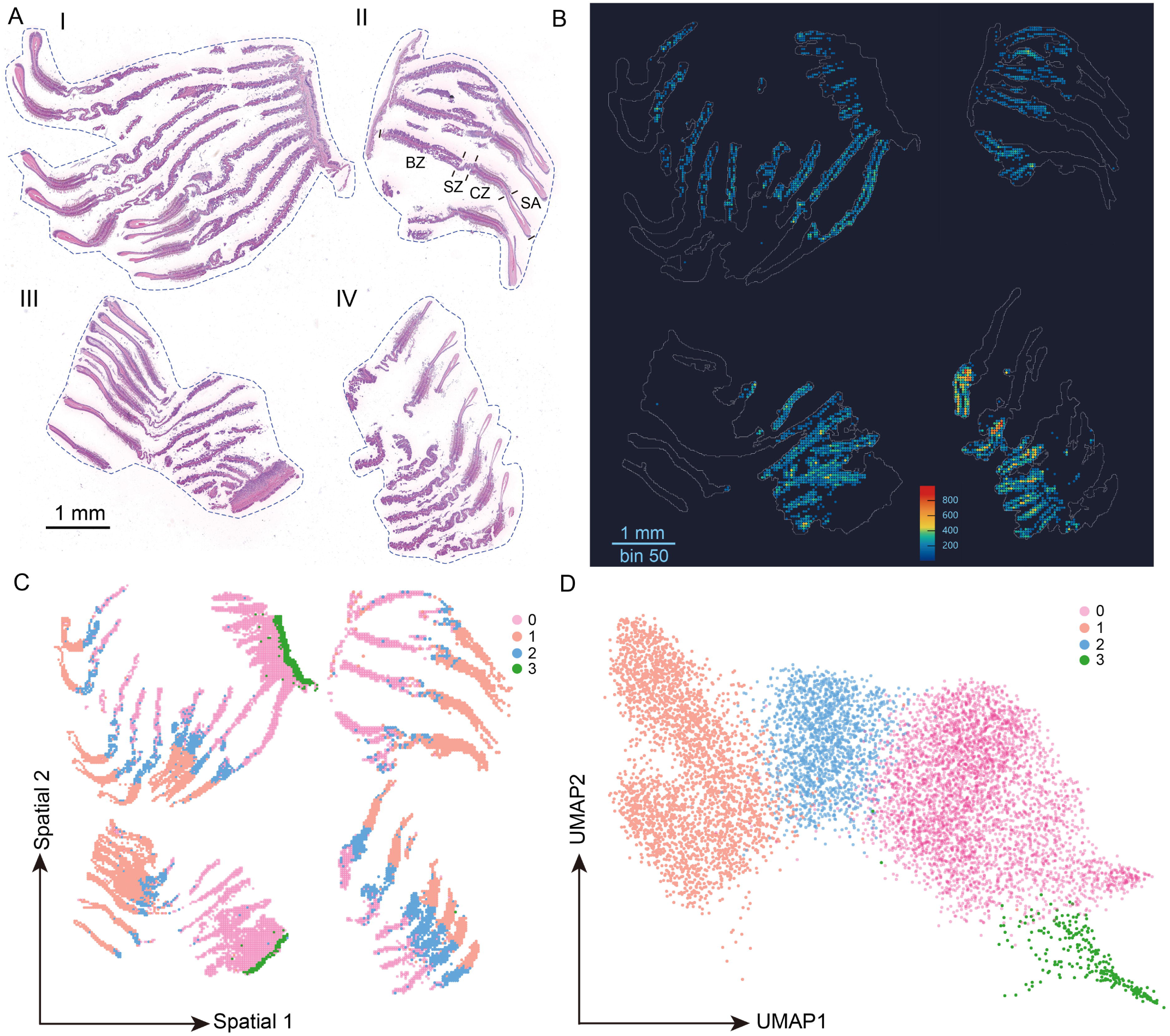
Spatial transcriptomics in *Alviniconcha marisindica* gill tissue. A, Semi-thin histological section stained with hematoxylin and eosin (HE) showing four independently prepared gill filaments. B, Distribution and abundance of 16S rRNA reads of *C*Sal in spatial transcriptomics, and the heatmap bar showing the relative 16S rRNA counts per bin. C, Spatial representation of clusters in the *A. marisindica* gill. D, Uniform manifold approximation and projection (UMAP) of clusters based on the gene expression level at the scale of bin 50 (i.e. 25µm).

For the host, a total of four clusters were presented in two dimensions using the method of Uniform Manifold Approximation and Projection (UMAP) of spatial transcriptome reads (Figure 4D). On the spatial scale, Cluster 0 is specially located in the BZ region, which corroborates with the locations of the symbionts; Cluster 1 exhibits the co-localizations in both CZ and SA regions, indicating similar functions of both cell types; Cluster 2 is located in SZ region (an area between CZ and BZ), the presence of this region has been known from other caenogastropods but has never been reported in *Alviniconcha*; Cluster 3 is located in the area that is likely related to the efferent branchial vessel (EBV) (Laming et al., 2020) (Figure 4C). Collectively, this distribution pattern matches well the anatomical structure of gill filaments (except for the Cluster 1), indicating the presence of different functional characteristics in each area.

We obtained the significantly highly expressed genes in each cluster, and GO enrichment was performed on these gene lists (Table S9, S10, S11 and S12 in Supplementary Table). For CZ and SA (Cluster 1), genes related to microtubule-based movement, motile cilium, and dynein complex were enriched. These genes are usually the key components of cilia cells, which support the functions of cilia movement for facilitating the water current (Figure S7, S8). For BZ (Cluster 0), we show the dynamically highly expressed genes related to chemosymbiosis (Figure 5A). First, under the scenario of the expanded gene family of the transmembrane protein SLCs, three SLC genes were found to be highly expressed in Cluster 0, including SLC4A, a transmembrane protein responsible for the bicarbonate/carbonate (key source for the symbiont carbon fixation) transport; SLC26A, a sulfate transporter; and SLC7A for arginine transportation. These SLCs are likely to be linked with the substance exchange (of bicarbonate/carbonate, sulfate, and arginine) between host and symbiont (Figure S9A-C).

**Figure 5.**
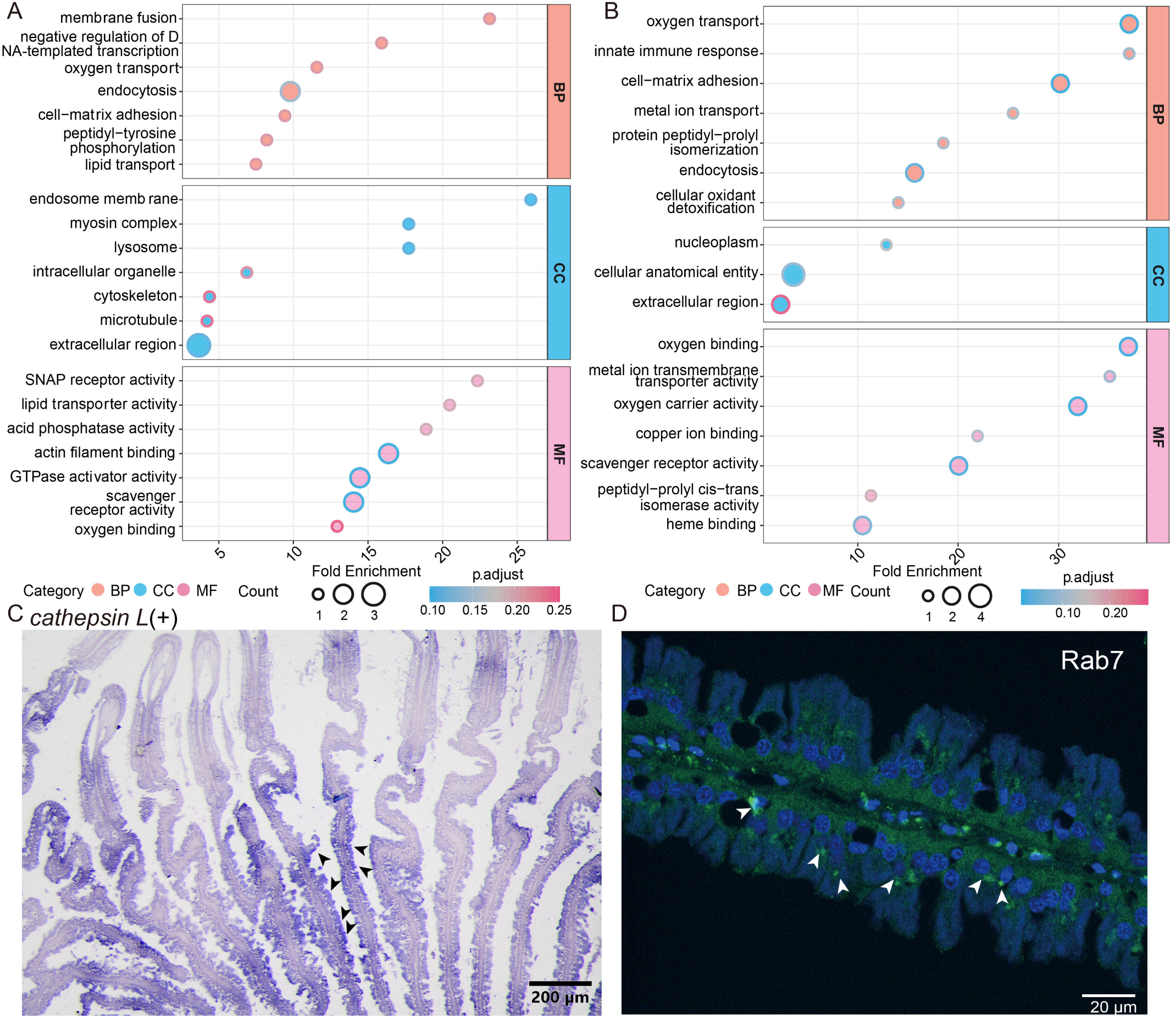
GO enrichment of cell type highly expressed genes. A and B, GO enrichment of significantly highly expressed genes (FDR < 0.05 and FC > 1.5) in the bacteriocyte zone and the secretory zone based on spatial transcriptomics, respectively. C, ISH of cathepsin L in *Alviniconcha marisindica* gill. Scale bar: 200µm. (**D**) IHC of Rab7 (green) and nucleus (blue) in *A. marisindica* gill. Scale bar: 20µm.

Secondly, we show that the symbiont digestion process can be akin to the endosome maturation; a similar process that is also be found in other chemosymbiosis species, such as *Riftia (Hinzke et al., 2019)* and *Bathymodiolus* mussels *(Chen et al., 2023b)*. Four genes related to endocytosis, receptor-type tyrosine-protein phosphatase beta (*PRTP-beta*) and two copies of deleted in malignant brain tumors 1 (*DMBT1*) (Figure S9D), were enriched, which may support the host obtaining symbionts horizontally via endocytosis (Sun et al., 2017). The symbionts are maintained in the endosome as revealed by the former and our TEM observations (Endow and Ohta, 1989) (Figure S10), and various genes were enriched to maintain the endosome stabilization, for instance, low-density lipoprotein receptor-related protein (*LRP*) gene and VAT-1 gene, which regulates vesicle transport (Kim et al., 2021). The lysosome, the final stage of the late-endosome, can help digest the symbionts (Chen et al., 2023b). We found several genes were involved, for instance, acid phosphatase type 7 (*ACP7*) gene (Figure S11A) and Prosaposin, two key marker genes in the lysosome (Chen et al., 2023b; Meyer et al., 2014). We also performed the IHC of Rab7, a maker protein of the late-stage endosome to lysosome (Wang et al., 2010), showing the higher protein expression level in the basal part of the bacteriocyte, in line with location of lysosome which is close to the location of the cell nucleus of BZ (Endow and Ohta, 1989) (Figure 5D, Figure S10, Figure S12). Genes related to various protein degradation or amino acid degradation were also found to be highly expressed in BZ, such as cathepsin L, digestive cysteine proteinase 1 (*DCP1*) and E3 ubiquitin-protein ligase rnf213-alpha (*RNF213*). The abovementioned genes may facilitate phagocytosis and symbiont digestion by the lysosome. To further validate this result, we performed ISH of cathepsin L, a key enzyme in lysosome for protein degradation (Ahn et al., 2010) while also serving as a crucial regulator of lysosomal function to control the residence of symbionts (Peyer et al., 2018), showing high gene expression level in the BZ (Figure5C, Figure S13, Figure S11B). In a nutshell, these gene and protein expressions highlighted the dominant “farming” strategy for nutrients in the BZ area.

For SZ (cluster 2), a symbiont-free area, genes involved in oxygen transport were enriched (Figure 5B). We call this zone the secretory zone (SZ), as it is positionally in agreement and probably homologous to the secretory zone known from gill filaments in other caenogastropods such as ampullairids (Rodriguez et al., 2019), where it mainly carries out mucus secretion. Here, we found three copies of mucin genes (mucin 2, mucin 4 and mucin-like) highly expressed in the SZ, corroborating the hypothesis that this zone is homologous to SZ in other caenogastropods. We also identified high oxygen transport in SZ; for instance, one myoglobin is highly expressed in SZ, and meanwhile, another globin, which can also bind to O_2_ (Yu et al., 2009), neuroglobin, is also highly expressed (Figure S11C and D). For the BZ, the gill filaments were filled with a dense population of symbionts, which can consume large amounts of O_2_ (Yu et al., 2009). Taken together, we propose a model of functional anatomy in *Alviniconcha* gill filaments involving three distinct functional zones: 1) In the CZ, oxygen supply is mediated through coordinated ciliary movements directed toward the BZ; 2) Within the BZ, molecular oxygen is primarily utilized by the outermost densely populated layer of symbiotic microorganisms; 3) The SZ serves as the critical site for the host’s respiration through oxygen binding and subsequent host-mediated oxygen uptake by respiratory proteins. This symbiont-free SZ is not known from the bathymodioline mussel gill filaments (Wang et al., 2024a). This is potentially due to 1) *Alviniconcha* lives closer to the vent edifice with a lower O_2_ concentration than *Bathymodiolus* mussels, requiring additional area in the gill for O_2_ binding; and 2) having a separate zone for host oxygen uptake may help the chemosymbionts thrive better in BZ, by reducing competition between the two parties in *Alviniconcha*.

## CONCLUSIONS

In summary, our multi-omics investigation of the *Alviniconcha* holobiont unravels a sophisticated interplay of genomic, proteomic, and physiological adaptations that underpin survival in the extreme and dynamic deep-sea hydrothermal vent ecosystems. The expansion and high expression of SLC families in *Alviniconcha* bacteriocytes highlight the host’s enhanced capacity for chemical/nutrient exchange with symbionts, while the metabolic complementarity in methionine biosynthesis underscores a critical dependency on symbiont-derived resources. The discovery of oxygen storage via myoglobin, rather than hemoglobin, resolves previous ambiguities and emphasizes a novel strategy to buffer fluctuating oxygen availability. Spatial transcriptomics further delineates functional partitioning within the gill, revealing bacteriocyte-mediated phagocytosis and lysosomal digestion as major routes for nutrient acquisition, alongside ciliary and secretory zones dedicated to oxygen transport and uptake. These adaptations, coupled with the symbionts’ streamlined genomes and reliance on host-derived metabolites, reflect a co-evolved holobiont system optimized for chemosynthetic mutualism. By integrating host-symbiont interactions across molecular and spatial scales, these results advance our understanding of how extremophile organisms exploit niche environments through synergistic partnerships, offering broader insights into the evolutionary innovation of chemosymbiosis across marine ecosystems.

## MATERIALS AND METHODS

### Sample collection, fixation, and Genome sequencing

*Alviniconcha marisindica* specimens were collected using the human-occupied vehicle (HOV) *Jiaolong* from the Wocan (6.36°N, 60.53°E) and Tianxiu (3°41′N, 63°50′E) vent fields on the Carlsberg Ridge, northwest Indian Ocean, during the research cruise DY38. Snails were flash-frozen in liquid nitrogen or dissected and preserved in RNAlater once recovered on-board the research vessel. For spatial transcriptome, FISH, IHC and ISH experiments, the *A. marisindica* gills were also fixed by 4% paraformaldehyde (PFA) at 4°C overnight and transferred to pure methanol at -30°C until use. *Alviniconcha adamantis* snails were collected by HOV *Shinkai 6500* from Suiyo Seamount vent field (28°34.2749′N, 140°38.6242′E) on the Izu-Ogasawara Arc at a depth of 1385 m on August 22, 2019 during R/V *Yokosuka* cruise YK19-10 (Figure S14). To avoid changes in gene expression during the recovery of snails from deep sea to the surface due to factors such as pressure and temperature, *A. adamantis* were fixed *in situ* in RNA-stabilizing solution following a published protocol (Sun et al., 2020; Yan et al., 2022). *A. marisindica* were immediately frozen in liquid nitrogen on board. The fixed individuals were dissected on-board the research vessel into different tissue types in RNA-stabilizing solution, and then kept at -80°C. Specimens for whole-genome sequencing were flash-frozen in liquid nitrogen and preserved in -80°C. For TEM the gill was fixed in 2.5% glutaraldehyde-seawater solution.

High-quality genomic extraction from mollusc can be difficult, as was also the case in our two *Alviniconcha* species (Sigwart et al., 2021). The genomic DNA was initially extracted using the sodium dodecyl sulfate (SDS)-based DNA extraction method and purified using the Genomic DNA Clean & Concentrator (gDCC-10) kit (ZYMO, CA, USA), a similar strategy was also applied to the Scaly-foot Snail genomic DNA extraction (Sun et al., 2020). Then, the purified DNA was sequenced on the R9.4.1 PromethION platform (Oxford Nanopore Technologies (ONT). The flash-frozen samples were subjected to Illumina library and Hi-C library preparing following our former protocol (Sun et al., 2020) and sequenced by Illumina HiSeq platform with PE 150 mode.

For ONT sequencing, the *A. marisindica* and *A. adamantis* raw *.fast5* data was basecalled into *.fastq* using Guppy v6.1.7 and Guppy v4.4.1 with the high accuracy (HAC) mode on GPU, respectively. The Illumina reads were obtained by removing bacterial contamination using Kraken2 (Wood et al., 2019). Low quality and sequencing adapter contaminated reads were trimmed by Trimmomatic v.0.39 (Bolger et al., 2014) with the settings of “LEADING=20 TRAILING=20 SLIDINGWINDOW = 4:20 MINLEN=50”. Calculations using three different kmer sizes, i.e. 17,19 and 21-mer, the heterozygosity of *A. adamantis* and *A. marisindica* was 0.68% and 1.67%. The total number of effective k-mers (total number of k-mers - total number of erroneous k-mers) divided by the number of homo-peak indicates that the estimated genome size of *A. adamantis* and *A. marisindica* was 640.58 Mb and 651.92Mb, respectively (Figure S15).

For mRNA sequences, tissue samples of mantle, foot, gill, radula sacs, neck, and operculum were dissected from two *A. marisindica* individuals from the Wocan vent field and another two from the Tianxiu vent field. For *A. adamantis*, mantle, gill, foot, digestive gland, and gonad of three *A. adamantis* individuals were sampled for transcriptome sequencing. Total RNA was extracted using Trizol reagent and sequenced separately on the Illumina NovaSeq 6000 platform with PE150 mode. At least three biological replicates were prepared for each sample. The details of host RNA-Seq data are summarized in Table S13 in Supplementary Table.

### Genome assembly

Seqtk v1.3 (https://github.com/lh3/seqtk) was used to remove short ONT sequences (<7 kb) for assembly using Flye v2.9 (Kolmogorov et al., 2019) with the settings of --nano-hq - scaffold. The assembled genome was polished two times with ONT reads. *A. marisindica* was assembled by Flye v2.9.1 (Kolmogorov et al., 2019) with the settings of “--no-alt-contigs” to remove all non-primary contigs from the assembly. Detailed benchmarking results of genome assemblies are shown in Table S14 in Supplementary Table.

The resultant genomes were larger than the estimated genome size by *k*-mer analysis. Therefore, purge-dups (Guan et al., 2020) was applied to further remove the heterozygous contigs. And the polca.sh from MaSuRCA v4.0.5 (Zimin et al., 2017) was further used to polish the genome with high-quality Illumina reads twice. BUSCO v5.2.2 (Seppey et al., 2019) was used to assess genome completeness after each step for selecting the optimal assembly result. To remove contaminated contigs, the details were shown in the supplementary materials (Figure S16, S17).

### Hi-C library preparation and sequencing

Additionally, Hi-C was used to anchor the primary genome assembly to chromosome level for the subsequent analyses. The chromatin was digested by the MboI restriction enzyme. Hi-C libraries were constructed and sequenced on Illumina NovaSeq platform with the read length of 150bp. The Illumina raw data were trimmed by Trimmomatic and the clean reads were aligned to the contigs by bowtie2 v2.4.5 (Langmead and Salzberg, 2012) with parameters of “--very-sensitive -L 30 --end-to-end”. To obtain the valid interaction paired reads, Hic-Pro v3.1.0 (Servant et al., 2015) was used to filter clean reads with the mapping quality of over 30. The resultant reads were aligned to primary genome assembly using Juicer v1.6 (Durand et al., 2016) and then genomic scaffolding the contigs into draft chromosomal assembly using 3D-DNA (Dudchenko et al., 2017) pipeline under default settings for diploid genomes. The resulted assembly was visualized and curated by Juicebox Assembly Tools (JBAT), with manually correction of misjoins, translocations, inversions, and chromosome boundaries. The run-asm-pipeline-post-review.sh pipeline was used to generate the corrected chromosomal assembly and chromosome heatmap. The chromosomes were sorted according to the length, and the longest chromosome was designated as Chromosome 01.

### Transcriptomic analysis

The RNA-seq reads of *A. marisindica* and *A. adamantis* were trimmed to ensure the reads’ quality and separately assembled using *de novo* and genome-guided modes in Trinity v2.13.2 (Haas et al., 2013) with the setting of “jaccard_clip” for the expected high gene density, respectively (Table S15 in Supplementary Table). Afterwards, the combination of two modes assembled transcripts were using CDHIT-EST v4.1.8 (Fu et al., 2012) to remove the high redundancy isoforms and further remove those short contigs which showed > 95% sequence similarities to the longer contigs. The transcripts deduced from the genome annotation were built as the index using salmon v1.8.0 (Patro et al., 2017) and the clean RNA-reads were quantified the index, and the quantification results were cross-sample normalized with *abundance_estimates_to_matrix.pl* from Trinity for downstream analysis with DESeq2. Four gill samples from *A. marisindica* were paired with those of other tissues from the same individual via the custom scripts using the condition of FDR < 0.05 to obtain highly expressed genes in gill (n = 4) (Table S6 in Supplementary Table).

### Genome annotation

To identify transposon elements (TEs) and other repeats, RepeatModeler v2.0.2a (Flynn et al., 2020) with default settings was used for simple repeats and transposable elements analyses, including the RMblast, to build a species-specific repeat library. RepeatMasker v4.1.0 (Flynn et al., 2020) searched, classified, and masked the repeats with the library and repeats in the Repbase database v2018.10.26. Therefore, repeat elements were identified using RepeatMasker twice with different repeat libraries with the hard-masked setting, one produced by searching the species-specific repeat library generated by RepeatModeler and the other produced by searching the public Repbase database. Finally, repeat elements were hard-masked for ease of annotation purposes.

For *ab initio* annotation, STAR v2.6.1a_08-27 (Dobin et al., 2013) was used for RNA-seq reads mapping to the genome. BRAKER v2.1.6 (Gabriel et al., 2021) was used to train AUGUSTUS v3.4.0 (Stanke et al., 2008) by the results of STAR and the hardmasked-genome from RepeatMasker. For homology annotation, the protein sequences of mollusk from the SWISS-PROT and NCBI database were mapped to the assembled genome using Exonerate v2.4.0 (Slater and Birney, 2005). For RNA-seq annotation, RNA-seq reads were trimmed by Trimmomatic and assembled using the *de novo* and genome-guided models in Trinity, respectively. Then, the combination of two assembled transcripts was mapped to the genome in the PASApipeline v2.5.2 (Haas et al., 2008) with gmap and blat as aligners.

The genomes were annotated using MAKER v3.01.04 (Cantarel et al., 2008) while setting the weight of evidence to improve the gene prediction accuracy. All gene evidence obtained above was collected and transferred to EVidenceModeler v1.1.1 (Haas et al., 2008; Haas et al., 2011) (EVM), which also can integrate evidence from different input and adjust the weight file for better results of gene prediction. The resultant .*gff3* files were updated using the PASA update pipeline twice to add UTR, and .fasta format file was converted by gffread v0.12.7 (Mihaela., 2020). To remove the isoform, TBtools v2.034 (Chen et al., 2023a) was used to pick the longest CDS from the gff3 file.

To functionally annotate the proteins in the *Alviniconcha* genomes, we used the protein sequences as the query to blast in the NCBI non-redundant database and SwissProt database using BLASTp with an E-value of 1e-5, respectively. The output files were uploaded to the Omicsbox v3.1.11 to obtain the GO annotation, and the clusterProfiler v4.0.5 (Yu et al., 2012) was used for GO enrichment analysis.

### Synteny analysis in *Alviniconcha*

The two *Alviniconcha* genomes were independently analyzed against with Mollusca Linkage groups (MLGs) from a previous study (Sigwart et al., 2025) through BLASTp search using diamond (Buchfink et al., 2015) with the threshold of ‘-evalue 0.001 -max_target_seqs 50000’. Strict one-to-one orthologous linkages were identified through a custom Python script implementing bidirectional best-hit criteria. Linkage group conservation was subsequently validated through pairwise dot plot analysis using the JCVI (Tang et al., 2008), followed by genome-wide macrosynteny visualization using the macrosyntR package (El Hilali and Copley, 2023).

### Gene family analysis

To examine the gene family expansion and contraction events, the 31 molluscan genomes were downloaded from NCBI and previous studies, with details shown in Table S4 in Supplementary Table). Cafe5 (Mendes et al., 2021) was used to determine the contraction and expansion of gene families, with the orthologous groups and genome phylogenetic tree (detailed Methods in SI, Figure S3) as inputs. Then, gene families with more than 100 gene copies were filtered out using *clade_and_size_filter.py* and the remaining gene families with *P* values lower than 0.01 were considered to be significantly expanded or contracted (Table S16 in Supplementary Table).

### Positive selection analysis

To identify the genes under positive selection in *Alviniconcha*, three models (two branch-site model BUSTED and aBSREL, a site model MEME) were used to compare them with a shallow water marine (*Monoplex corrugatus*) and a freshwater snail (*Marisa cornuarietis*). Mafft (Katoh and Standley, 2013) was used to align amino acid sequences, and amino acid alignment further guided the alignment of coding DNA sequences (CDS) in pal2nal (Suyama et al., 2006). On the basis of the phylogenetic tree, the CDS alignments of these genes were used to detect the positive selection pressure using BUSTED, aBSREL, and MEME models implemented in Hyphy (Kosakovsky Pond et al., 2019).

### Proteomics characterization

Gill samples from three individuals of *A. marisindica* were used in the proteomic analysis. First, the lysis buffer (8M urea and 1M HEPES pH = 7.4) was used to lyse the samples into a homogenate and then centrifuged to remove the cell debris at 14,000g for 10 min at 4 °C to collect the supernatant that contained proteins. The supernatant was purified, checked with SDS-PAGE, and digested with Trypsin (Promega) to gain the peptides. The peptides were passed to LC-MS/MS detection via a tandem mass spectrometer Q-Exactive HF X (Thermo Fisher Scientific, San Jose, CA). The MS/MS data was identified and quantified via MaxQuant v1.5.3.30 (Sinitcyn et al., 2021), searching two reference (host and symbiont) databases plus their reversed sequences, which served as the “decoy” database, respectively. Two searched reference databases include the *A. marisindica* translated protein database from the genome, its symbiont protein database from a previous study (Yang et al., 2020) and those reversed proteins that served as the decoy, correspondingly. The protein-level FDR < 0.01 and PSM-level FDR < 0.01 were the selected conditions to obtain the significantly identified proteins.

### Spatial transcriptomics

We applied the stereo-seq technique (BGI, China) for the spatial transcriptome sequencing following our formerly established protocol (Li et al., 2025). Briefly, tissue sections were adhered to the surface of Stereo-seq N FFPE Chip (BGI, China) and incubated at 37 °C for 3 mins. The sections were then fixed in methanol and incubated at -20 °C for 40 mins prior to stereo-seq library preparation. Then, sections were stained with Hematoxylin and Eosin (HE) and imaged using a SDPTOP HS6 for capture.

The section was washed using 0.1 x SSC buffer with 0.05U/ml RNase inhibitor, and then permeabilized using 0.1% pepsin in 0.01 M HCl buffer. The sections were incubated at 37 °C for 5 mins and washed again with 0.1 x SSC buffer containing 0.05 U/ml RNase inhibitor. RNA was captured by DNA nanoballs (DNB) and reverse-transcribed overnight at 42 °C using SuperScript II according to the standard method. After reverse transcription, sections were washed twice with 0.1 x SSC buffer and digested at 55 °C with tissue removal buffer (10 mM Tris-HCl, 25 mM EDTA, 100 mM NaCl, 0.5% SDS) for 10 mins. The Chips were treated with cDNA Release Mix overnight at 55 °C. The VAHTSTM DNA Clean Beads (0.8x) was used to purify cDNA.

The resultant cDNA was amplified with KAPA HiFi Hotstart Ready Mix (Roche, KK2602) and the concentration of PCR products was determined using Qubit dsDNA Assay Kit. DNA was fragmented with TnT transposase at 55 °C for 10 mins, and then the reaction was stopped by adding 0.02% SDS and incubating at 37 °C for 5 mins. The fragmented DNA was amplified as follows: 25 μl of the fragmentation product, 1x KAPA HiFi Hot-start Ready Mix, along with 0.3 mM of each Stereo-seq-Library-F and Stereo-seq-Library-R primers, were mixed with nuclease-free H2O to a total volume of 100 μl. The PCR products were purified using AMPure XP Beads to generate DNB, and sequenced on the MGI DNBSEQ-Tx sequencer (BGI) with approximately 600Gb of raw reads.

Reads 1 contains CID (25bp) and MID (6bp), while Read 2 consists of the cDNA sequences. CID sequences in Read 1 were first mapped to the *in situ* chip coordinates obtained from the first sequencing round, followed by 1 base mismatch to account for sequencing and PCR errors. Reads containing MID sequences with either N bases or more than 2 bases with a quality score below 10 were filtered out. The CID and MID information associated with each read was appended to the read headers. Retained reads were aligned to the *A. marisindica* genome delimitated by the .gff annotation file using STAR (Dobin et al., 2013), and only reads with MAPQ > 10 were counted and annotated to their corresponding genes. UMI with identical CID and gene locus were collapsed, allowing for 1 mismatch to correct for sequencing and PCR errors. Finally, this data was used to generate a CID-containing expression profile matrix. The entire workflow was implemented in the publicly available SAW pipeline (https://github.com/BGIResearch/SAW).

The expression profile matrix was divided into non-overlapping 8,919 bins, each covering an area of 50 x 50 DNB. The resulting cells were processed using Scanpy (Wolf et al., 2018) and Stereopy (https://github.com/STOmics/Stereopy), followed by normalization, marker gene selection, PCA dimensionality reduction, and clustering.

### *In situ* hybridization (ISH), Fluorescence *in situ* hybridization (FISH) and Immunohistochemistry (IHC) Samples and probe synthesis

According to the manufacturer’s instructions, total RNA was extracted using an Eastep ® Super Total RNA Extraction Kit (Promega, USA) and purified using the RNA purification and recovery Kit (Genstone, China). The first strand of cDNA from the gill RNA was synthesized using a GoScript^TM^ Reverse Transcriptase kit (Promega, USA) according to the manufacturer’s instruction and used for PCR amplification (These primers were shown in Table S17 in Supplementary Table). Then, the PCR products were cloned into the pEASY-T3 vector (TransGen, China), and the clone plasmids were purified using the TIANprep Mini Plasmid Kit (TIANGEN, China) and sequenced using the Sanger method. The plasmids were digested by SphI restriction enzyme (Takara, Japan) for the subsequent probe synthesis. By *in vitro* transcription using the linear plasmids as the template, anti-sense and sense RNA probes were synthesized with SP6 and T7 RNA polymerase (Thermo Scientific, Germany), respectively, using 2µl 10x DIG RNA Labeling Mix (Roche, Switzerland) according to the manufacturer’s instructions. After the synthesis of RNA probes, the linear plasmids were digested by DNase I at 37 °C for 20 min. The RNA probes were finally dissolved in DEPC-treated water and preserved at -80 °C for further experiments.

## ISH

The gill was dehydrated through an ethanol series and embedded in paraffin (Leica, Germany). The 5µm sections were cut with a rotary automatic microtome (Leica, Germany). The 5µm paraffin sections on covering slides were incubated at 37 °C overnight.

Afterwards, the sections were rehydrated through xylene, ethanol series (100%, 95%, 85%, 75%) and washed with 0.1% Tween in phosphate-buffered saline (PBS) for 10 min. The sections were washed with PBST and digested with Proteinase K (10µg/ml) in incubation box at 37 °C for 15 min, stopped with PBST for 10 min, and fixed in 4% PFA/PBS for 20 min at room temperature. The sections were subsequently subjected to 200µl of pre-hybridization buffer (4.78mM Citric acid PH 6.0, 2.5% deionized formamide, 2.5% SSC, 0.05 µg/mL heparin, 0.5 µg/mL torula yeast RNA, and 1‰ Tween) in incubation box at 60 °C for 1 h. The pre-hybridization buffer was replaced by the hybridization buffer with a final concentration of 1µg/ml DIG-labeled probes at 60 °C overnight. After hybridization, the sections were washed three times in 50% Formamide/2x SSCT at 60 °C for 30 min, twice in 0.2x SSCT at 60 °C for 30 min. Later, the sections were washed twice in PBST for 30 min at room temperature and then incubated with 0.5% blocking reagent for 30 min at RT. The AP-conjugated anti-DIG antibody in the blocking reagent (1:2000) was added to the samples at 4 °C overnight. Then, the PBST was used four times for 30 min to remove excess the antibody and the sections were washed in TNMT buffer (100mM Tris PH9.5, 50mM MgCl_2_, 100mM NaCl, 1‰ Tween) for 5 min. The alkaline phosphatase activity was detected with BM purple AP substrate. After the color development, the sections were washed with PBST twice for 5 min, immediately mounted in 70% glycerol with coverslips and observed under a dissecting binocular (Olympus SZX16, Japan).

## FISH

The sections (4µm) were carried out as described above, and the sections were hybridized with 50ng/ml EUB338-Cy5 probe to target 16S rRNA of symbionts in a hybridization buffer (0.9M NaCl, 0.02 M Tris-HCl, 0.01% SDS and 20% formamide) at 46 °C for 1h. After washing three times with a wash buffer (0.1 M NaCl, 0.02 M Tris-HCl, 0.01% sodium dodecyl sulphate, and 5 mM EDTA) at 48 °C for 15 min, the 4′,6-diamidino-2-phenylindole (DAPI, Solarbio, China) and Alexa Fluor 488 Conjugate Concanavalin-A (Invitrogen, CA, USA) were labelled with sections to target nucleotide acids and cell membrane, respectively. Then, the sections were mounted on a Prolong glass antifade mounting medium (Invitrogen, CA, USA) and then imaged under a confocal microscope (Nikon Eclipse T*i*2, Japan).

## IHC

For IHC analysis, the sections (5µm) were carried out as described above, and the sections were incubated with 1: 200 diluted anti-Rab-7 rabbit antibody (Abcam) with Block buffer (2% BSA and 2% sheep serum) at 4 °C for overnight, and then incubated with 1:1000 diluted Alexa Fluor 488 labeled goat anti-rabbit secondary antibody (Invitrogen) with Block buffer at RT for 2.5 h. Then, the sections were stained with DAPI and observed and imaged under a confocal microscope (Leica dmi8, Germany).

## Compliance and ethics

The authors declare that they have no conflict of interest.

## Data availability

All data needed to evaluate the conclusions in the paper are present in the paper and/or the Supplementary Materials. All sequencing data, including the raw Illumina, ONT, HiC, transcriptome sequencing and stereo-seq sequencing, were deposited on NCBI under the BioProject number of PRJNA1245653. This study also used data with NCBI number of GCA_020525365.1. The genome assemblies were deposited on the figshare (https://doi.org/10.6084/m9.figshare.28670054.v2). The raw mass spectrometry data were deposited in the ProteomeXchange Consortium via PRIDE with the accession number of PXD062809.

## Acknowledgement

This work was financially supported by National Key Research and Development Program of China (2024YFC2816100), the Science and Technology Innovation Project of Laoshan Laboratory (LSKJ202203104, 2022QNLM030004-2 and project no. LSKJ202203206), the Natural Science Foundation of Shandong Province (ZR2023JQ014), and the Young Taishan Scholars Program of Shandong Province (tsqn202103036). The authors would like to thank the captain and crew members of *Xiangyanghong 09* and *Shenhaiyihao*, and pilots of *Jiaolong* during the cruise of DY38 and DY 72, as well as the captain and crew of R/V *Yokosuka* and the HOV *Shinkai 6500* team during research cruise YK19-10. This is a GACHINKO Cruise Episode I (YK19-10) output; Ken Takai (JAMSTEC) is thanked for his diligent efforts in leading the cruise and the GACHI-participants for their help in sample preparation. We thank the ‘GACHI-Fighters’ (undergraduate students) of GACHINKO Cruise Episode I in helping with sample sorting on-board R/V *Yokosuka*. Katsuyuki Uematsu (Marine Works Japan Ltd.) is gratefully acknowledged for his great help in transmission electron microscopy.

## Notes

### Competing Interest Statement

The authors have declared no competing interest.

