## Supplementary Materials for "Multiple-omics analyses of the *Alviniconcha* holobiont reveal multi-faceted adaptations to deep-sea hydrothermal vents"

Molecular adaptation of two hot-vent endemic *Alviniconcha* gastropods to the chemosymbiosis as revealed by multiple omics studies

Wang *et al.*

**This PDF file includes:**

Supplementary Text

Figure S1 to S17

References (1 to 33)

**Other Supplementary Material for this manuscript includes the following:**

Supplementary Tables

Supplementary Text

**Genome survey**

The *K*-mer frequency with a *K*-mer size of 17, 19 and 21 was counted by Jellyfish v.2.3.0 (Marçais and Kingsford, 2011), and the resultant *K*-mer histograms were submitted to GenomeScope 2.0 (Ranallo-Benavidez et al., 2020) to get the genome size and heterozygosity. Calculations using three different kmer sizes, i.e. 17,19 and 21-mer, the heterozygosity of *A. adamantis* and *A. marisindica* was 0.68% and 1.67%. The total number of effective *K*-mers (total number of k-mers - total number of erroneous *K*-mers) divided by the number of homo-peak indicates that the estimated genome size of *A. adamantis* and *A. marisindica* was 640.58 Mb and 651.92 Mb. *K*-mer plot showing the distribution of *K*-mer copy number (KCN) at 19 for the *Alviniconcha* genome. To better visualize the distribution of KCN, only values between 0 and 120 were shown in Figure S15.

**Genome assembly and quality assessment of assembled genomes**

We performed a number of benchmarking studies to obtain a high-quality genome with the best continuity and completeness. Seqtk v1.3 was used to remove short sequences for assembly, which are less than the set length. Shasta (Shafin et al., 2020) and Raven5 (Vaser and Šikić, 2021) were designed for ONT data assembly, and Flye was recommended for assembling genomes with less heterozygosity. The assembled genome results by setting the parameters of different assemblers are shown in Table S14 in Supplementary Tables. Considering the overall, N50, NG50 and BUSCO are frequently used to evaluate the completeness of genome assembly. Based on Table S14, the best assembly of *A. adamantis* came from the assembly using Flye v2.9 (Kolmogorov et al., 2019) on the ONT reads longer than 7kb. The assembled genomes were polished two times by Flye v2.9. The resultant genome was larger than the estimated genome size by k-mer analysis. Therefore, purge-dup (Guan et al., 2020) was applied to further remove the heterozygous contigs. And the polca.sh from MaSuRCA v4.0.5 (Zimin et al., 2017) was used two times for polishing the genome with Illumina reads. BUSCO v5.2.2 (Seppey et al., 2019) was used to assess genome completeness after each step for selecting the optimal assembly result.

To remove contamination contigs, minimap2 v2.2.2 (Li, 2018) and samtools v1.138 (Danecek et al., 2021) were used to map the genome using ONT raw reads. Blobtools (Challis et al., 2020) was used to check the microbe-contaminations by applying the following three steps: 1) aligning assembled contigs against NCBI nt database by BLASTn with an E-value cutoff of 1e-5; 2) calculating the contig sequencing depth generated by minimap2; and 3) calculating the the contig average GC content (Figure S16 and Figure S17). To further confirm the decontamination results from Blobotools, the contig sequences were also examined by Megan v6.23.4 (Huson et al., 2016). The BUSCO score remains unchanged in this step. In the end, the size of the *A. adamantis* genome was 794Mb, and the BUSCO score was 95.0% complete single copy gene models (C), 0.6% complete duplicate gene models (D), 2.8% fragmented gene models (F) and 2.2% missing gene models (M).

Likewise, *A. marisindica* was directly assembled by Flye v2.9.1 with the settings of “--no-alt-contigs” to remove all non-primary contigs from the assembly (Table S14 in Supplementary Tables). Then, referring to *A. adamantis* assembly and deduplication pipeline, the genome size of *A. marisindica* was 841Mb, with a total of 1611 contigs and a scaffold of N50 of 2.09Mb. The BUSCO result of *A. marisindica* revealed 94.0% C, 0.9%D, 3.6% F and 2.4%M.

**Phylogenomic analysis of the *Alviniconcha***

To determine the phylogenetic positions of the *Alviniconcha* in Gastropoda, transcriptomes within Gastropoda were downloaded from the NCBI SRA database, and the genomes were downloaded from either NCBI genome database or the data link provided by the corresponding papers, with details shown in Table S3 in Supplementary Tables. The complete table of specimens, including information on the origin, BUSCO score and number of proteins, as well as the details and taxonomic information, are provided as Table S3 in Supplementary Tables.

The raw data were *de novo* assembled by Trinity v2.13.2 or v2.14.2 after removing the adapters based on data type by Trimmomatic. Afterward, Transdecoder v5.5.0 (Douglas, 2018) was used to convert transcripts to proteins according to the open reading frame, and then redundant transcript isoforms were removed by CD-HIT-EST (Fu et al., 2012) with a strict threshold (-c 0.8 or 0.75), which can obtain the lowest “D” BUSCO score.

The phylogenomic pipeline followed a previously published protocol (Li et al., 2025; Liu et al., 2023). We searched for orthologues in the selected taxa by Orthofinder v2.5.4 (Emms and Kelly, 2019), and then 50% occupancy was set before deleting the sequences shorter than 100 amino acids. The redundant sequences were removed by uniqHaplo.pl, and the remaining sequences were aligned by MAFFT v7.490 (Katoh and Standley, 2013). HmmCleaner v0.180750 (Di Franco et al., 2019) was searched to remove mis-aligned sequences and BMGE v.1.12 (Criscuolo and Gribaldo, 2010) was used to trim ambiguously alignment, followed by making a fast maximum-likelihood (ML) tree for each OG with FastTree2 (Price et al., 2010). PhyloPyPrunner v1.2.4 (<https://pypi.org/project/phylo-pypruner>) was used to identify the orthologs based on the FastTree results to generate an original matrix containing 3127 single-copy OGs. The origin matrix was constructed as an IQ-TREE2 (Minh et al., 2020) with “MFP” mode. GenesortR (Nicolás., 2021) was performed to sort and subsample the OG datasets by calculating seven gene properties to quantify phylogenetic usefulness for downstream analyses. Each OG was used to construct an MFP tree and the resultant trees were merged as the input file in genesortR. By setting the desired number of genes in the final datasets, we obtained three matrices, including 600 genes matrix, 800 genes matrix, and 1200 genes matrix. Except for the original matrix, other matrices were separately performed using IQ-TREE with “-m MFP” and “-m LG+C60” for phylogenetic analysis. Radom 300 genes from the original matrix were performed using Phylobayes MPI v.1.8c (Lartillot et al., 2013) with the model of CAT+GTR. Afterward, we chose the most supported topology from all trees generated above as the final tree.

**Time-tree analysis**

MCMCTree in PAML v4.10.5 (Yang, 2007) was used to estimate the divergent time of *Alviniconcha* from other lineages, which selects the final tree as the reference tree topology. We set fossil calibrations based on our previous study (Sun et al., 2021) : 1) a soft constraint of 520.5 Ma and 530 Ma for the origin of Bivalvia; 2) the split of *Aplysia* and *Biomphalaria* at a hard minimum bound of 168.6 Ma and a soft maximum bound of 473.4 Ma ; 3) the split between Caenogastropoda and Heterobranchia at a hard minimum bound of 390 Ma; 4) a hard constraint of 470.2 Ma and a soft constraint of 531.5 Ma for the first appearance of Gastropoda; 5) a hard minimum bound of 532 Mya and a soft maximum of Ma for the first appearance of Mollusca; 6) the split between the old world ampullariids and the new world ampullariids at a hard maximum bound of 150 Ma; 6) the split between Neogastropoda and other Caenogastropoda at a hard minimum bound of 144 Ma (Osca et al., 2014); 7) a time for root at 549 Ma. The fossil-phylogenetic tree and the protein sequence alignments were imported into co*deml* to roughly estimate the substitution rate. According to the overall substitution rate, the rgene gamma of *MCMCTree.ctl* was set as (1, 17.9). The final run of MCMCTree with the settings “burn-in = 10000 sampfreq = 100 nsample = 200000”. The resultant tree was visualized by FigTree v1.4.4 (<https://github.com/rambaut/figtree/>) (Figure S2).

In our analysis, all methods and matrices congruently placed the two *Alviniconcha* species in the subclass Caenogastropoda with maximum support, among other taxa currently assigned to the order Littorinimorpha (Figure S2). Like other previous phylogenetic studies of Caenogastropoda, our tree also revealed a paraphyletic of Littorinimorpha (Wang et al., 2024). Fossil-calibrated analysis based on the current phylogeny tree with 7 calibration points suggested that the genus *Alviniconcha* diverged from other gastropods at around 114.81 million years (Ma) ago (95% HPD: 76.68 – 156.91), and the two *Alviniconcha* species diverged approximately 15 Ma (95% HPD: 7.32 – 25.32). The divergent time between the two species recovered is slightly later than a previous study (Corinna et al., 2020) but overall overlap considering the 95% HPD.

**Identification of Myoglobin**

We used protein sequences that were downloaded from uniprot database and the previous study (Hoffmann et al., 2010). Afterward, the 3D protein structure files (.pdb) of each protein were constructed using colab-fold v1.5.5 (Mirdita et al., 2022) with the settings of “--num-relax 1 --num-recycle 3 --model-type alphafold2_ptm”. The inter-protein phylogeny was constructed based on the protein pdb files via FoldMason (Gilchrist et al., 2024), this result also supports the myoglobin existed in *Alviniconcha* due to the myoglobin with myoglobin gastropoda forming a clade (Figure S4A). Figure S4B showed the sense myoglobin probe (negative result) in *Alviniconcha marisindica* gill section.

**Transmission Electron Microscopy**

The gill tissue of *A. adamantis* was dehydrated through a gradient acetone series, and then embedded in Epon resin (Sigma-Aldrich). Ultrathin (70 nm) sections were cut using an ultramicrotome (Reichert Ultracut S, Leica). The sections were stained in 2% aqueous uranyl acetate with lead stain solution (0.3% lead acetate and 0.3% lead nitrate, Sigma-Aldrich). TEM observations were carried out using a Tecnai 20 Transmission Electron Microscopy (FEI), at an acceleration voltage of 120 kV.

Transmission electron micrographs (TEM) of the *A. adamantis* gill confirmed that the bacteriocytes have microvilli on their surfaces, which may help O_2_ exchange and also obtain symbionts horizontally from the environment (Figure S10A). The slender rod-shaped symbionts were densely packed near the external surface of the bacteriocytes, while lysosomes were localized in the inner part of the bacteriocytes. The cell membrane of the bacteriocytes were not enclosed, confirming the *Alviniconcha* symbionts are not completely internalized in the host cell (Endow and Ohta, 1988). This pattern is comparable with the FISH results, and a similar pattern was previously also observed in another congener, *Alviniconcha hessleri* (Endow and Ohta, 1988).

**Spatial transcriptome**

Under bin50, we selected eight genes (*SLC4A4*: scaffold_410_1.305, *SLC7A2*: scaffold_556_1.50.1.6348841c, *SLC26A6*: contig_3803_1.18, *DMBT1*: scaffold_702_1.121 and scaffold_702_1.122 , *myoglobin*: contig_3803_1.35, *cathepsin L*: contig_441_1.36, and *neuroglobin*: contig_2193_1.55 and *acid phosphatase type 7* (*ACP7*): contig_2067_1.302) that were highly expressed in BZ region and also mentioned in the maintext to show the distribution in *Alviniconcha marisindica* gill filaments (Figure S10-11).

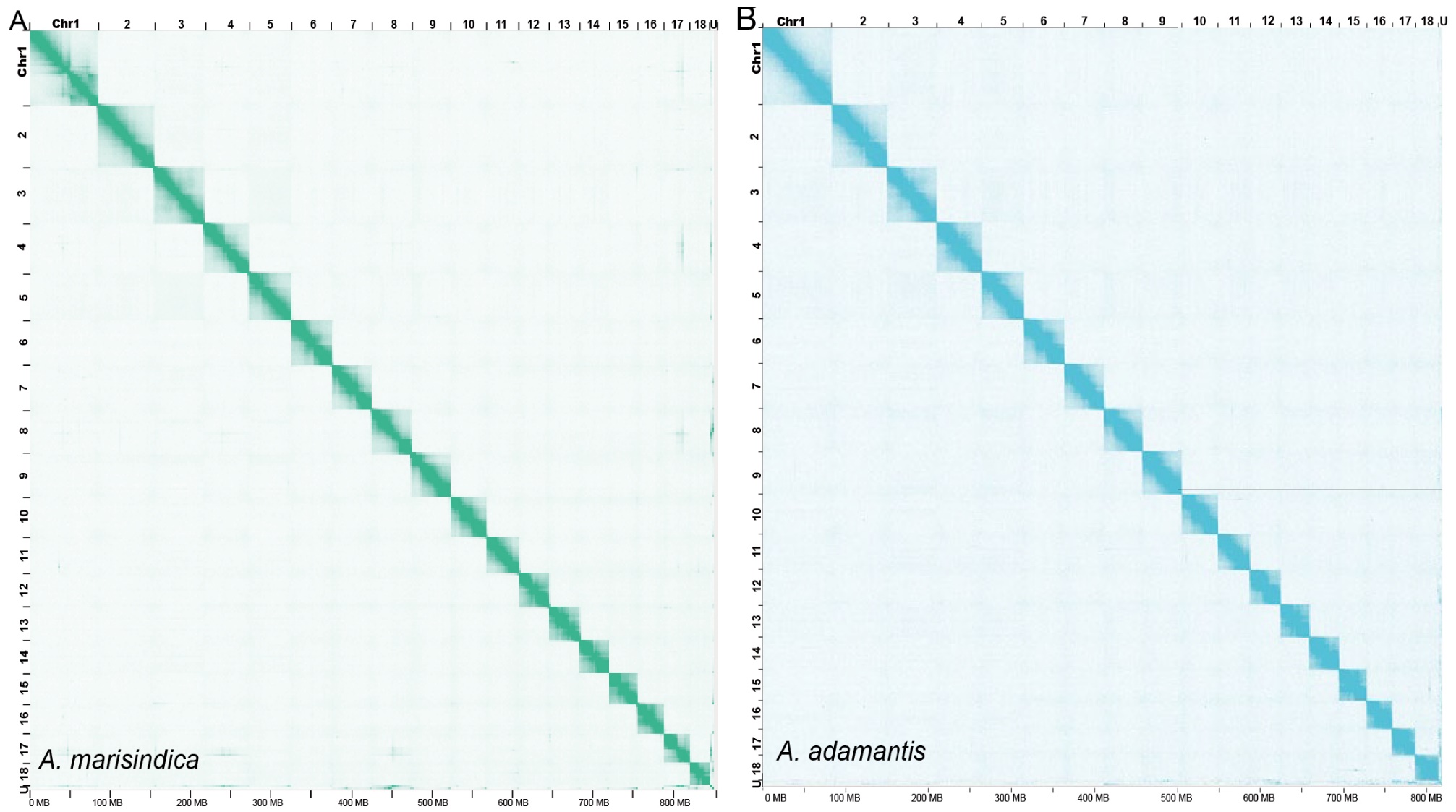

**Figure S1.** **The Hi-C interaction** **map showing the orientation of chromosomes in two *Alviniconcha* genomes.** (A) Genome-wide Hi-C interactive heatmap of *A. marisindica*. (B) Genome-wide Hi-C interactive heatmap of *A. adamantis*.

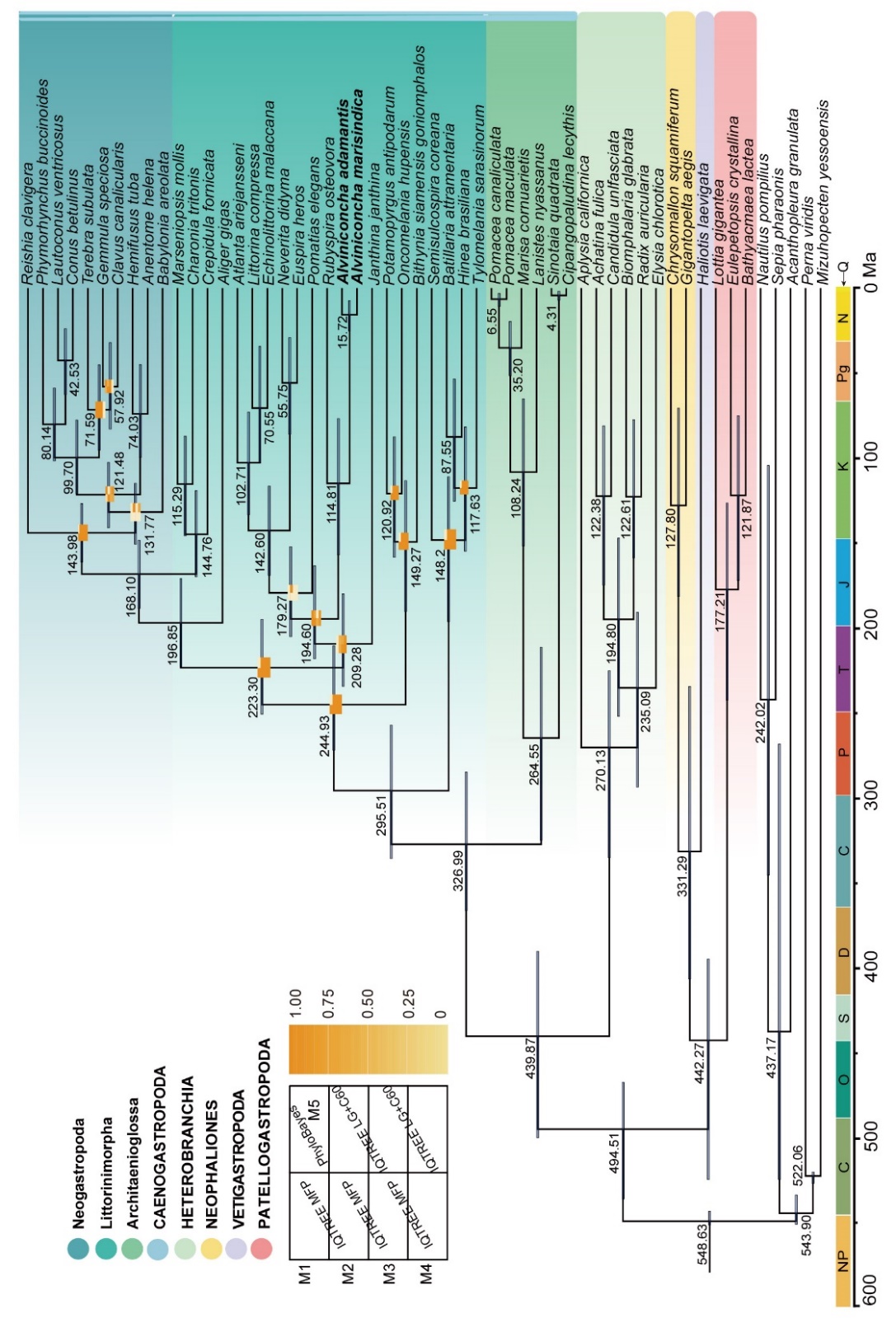
**Figure S2. Phylogeny of Gastropoda based on a phylogenomic approach using different softwares and models.** All branches are full supported, except the heatmap branches. The heatmap from light orange to dark orange indicates the support value from 0 to 100. N: Neogene; Pg: Palaeogene; K: Cretaceous; J: Jurassic; T: Triassic; P: Permian; C: Carboniferous; D: Devonian; S: Silurian; O: Ordovician; C: Cambrian.M1: all gene matrix. M2: 1200 gene matrix; M3: 800 gene matrix; M4: 500 gene matrix; M5: 300 gene matrix. The times of speciation are marked on the branches, and the range of light purple bars indicates the 95% confidence interval of the divergence time.

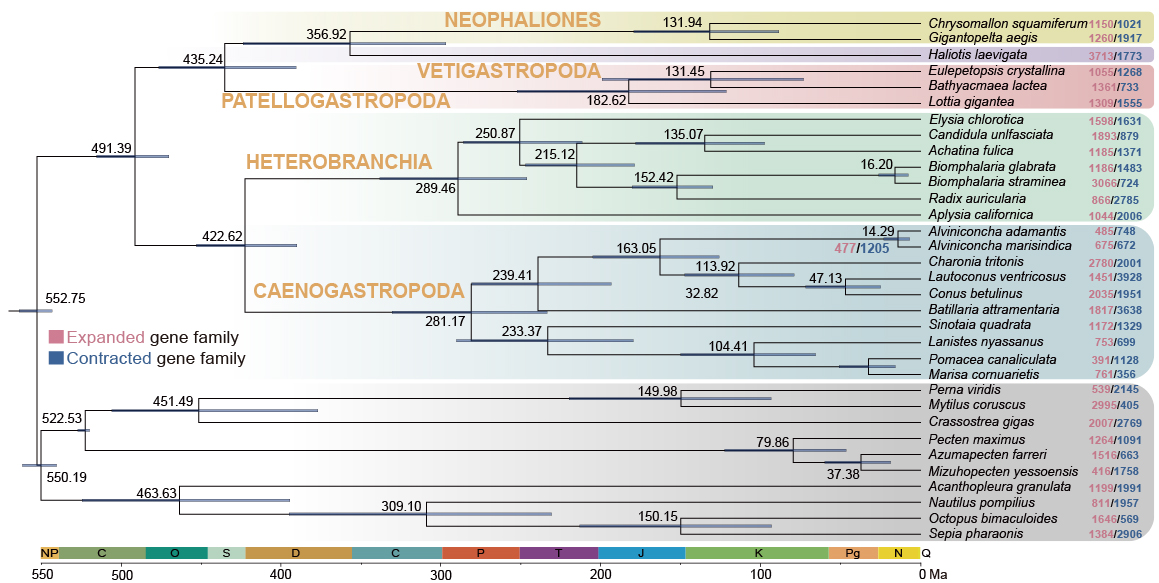

**Figure S3. Gene family expansion/contraction analysis for 33 selected species.** The number in pink represents the expanded gene families, and the number in blue represents the contracted gene families.

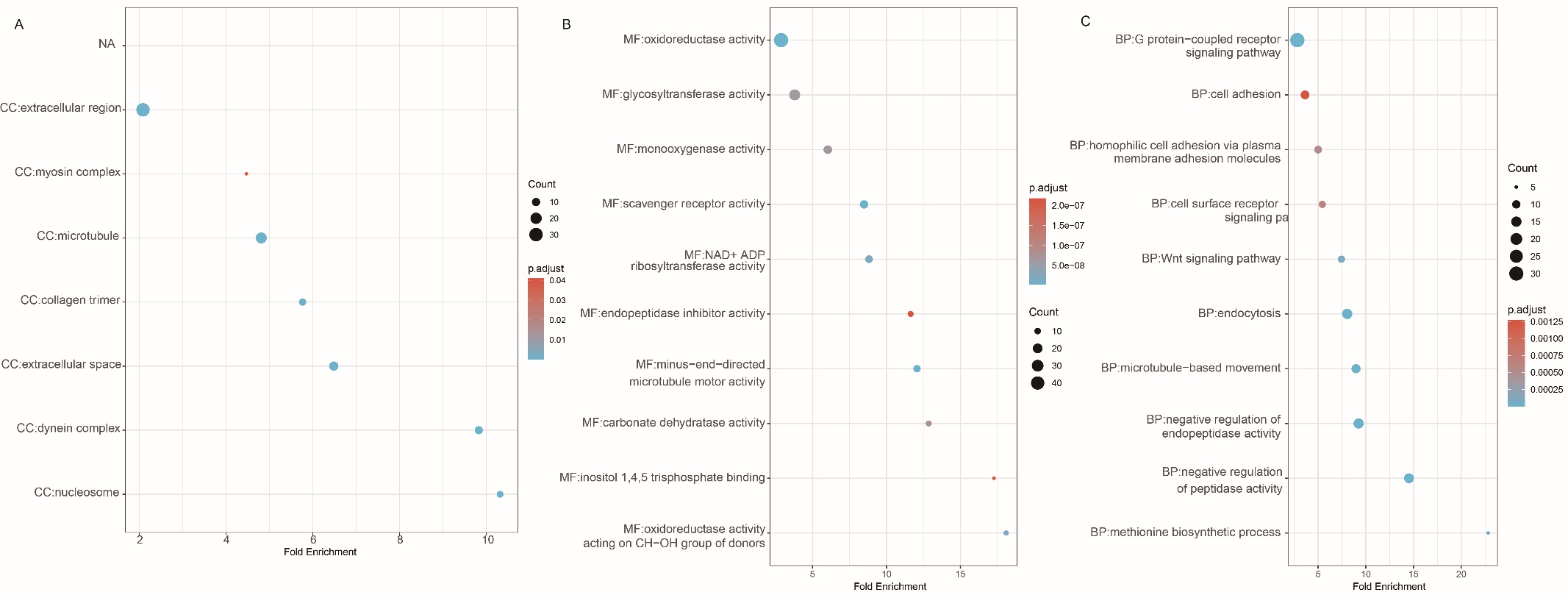

**Figure S4. GO Enrichment analyses of significant contracted gene families in two *Alviniconcha*.** GO terms that exhibit statistically significant differences are shown in the graph. (a) GO cellular component terms enriched in contracted genes;(b) GO molecular function terms enriched in contracted genes; (c) GO biological process terms enriched in contracted genes.

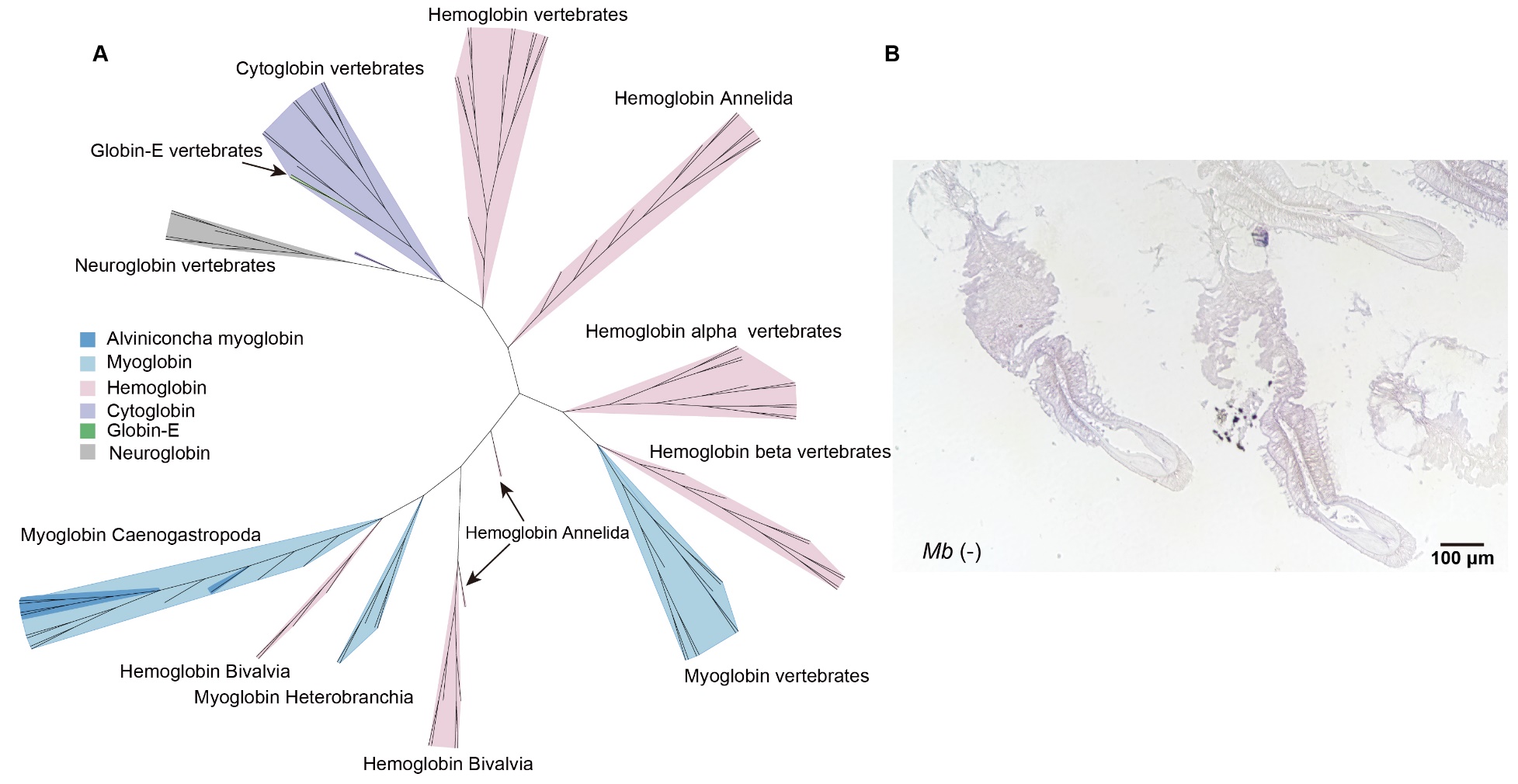

**Figure S5. Identify of myglobin and expression localize.** (A) The unrooted Phylogeny tree was constructed with 3D protein structure of *Alviniconcha* myoglobin and other typical globin proteins via FoldMason. (B) Negative control of *in situ* hybridization to detect myoglobin in the *Alviniconcha marisindica* gill filaments. Scale bar: 100μm.

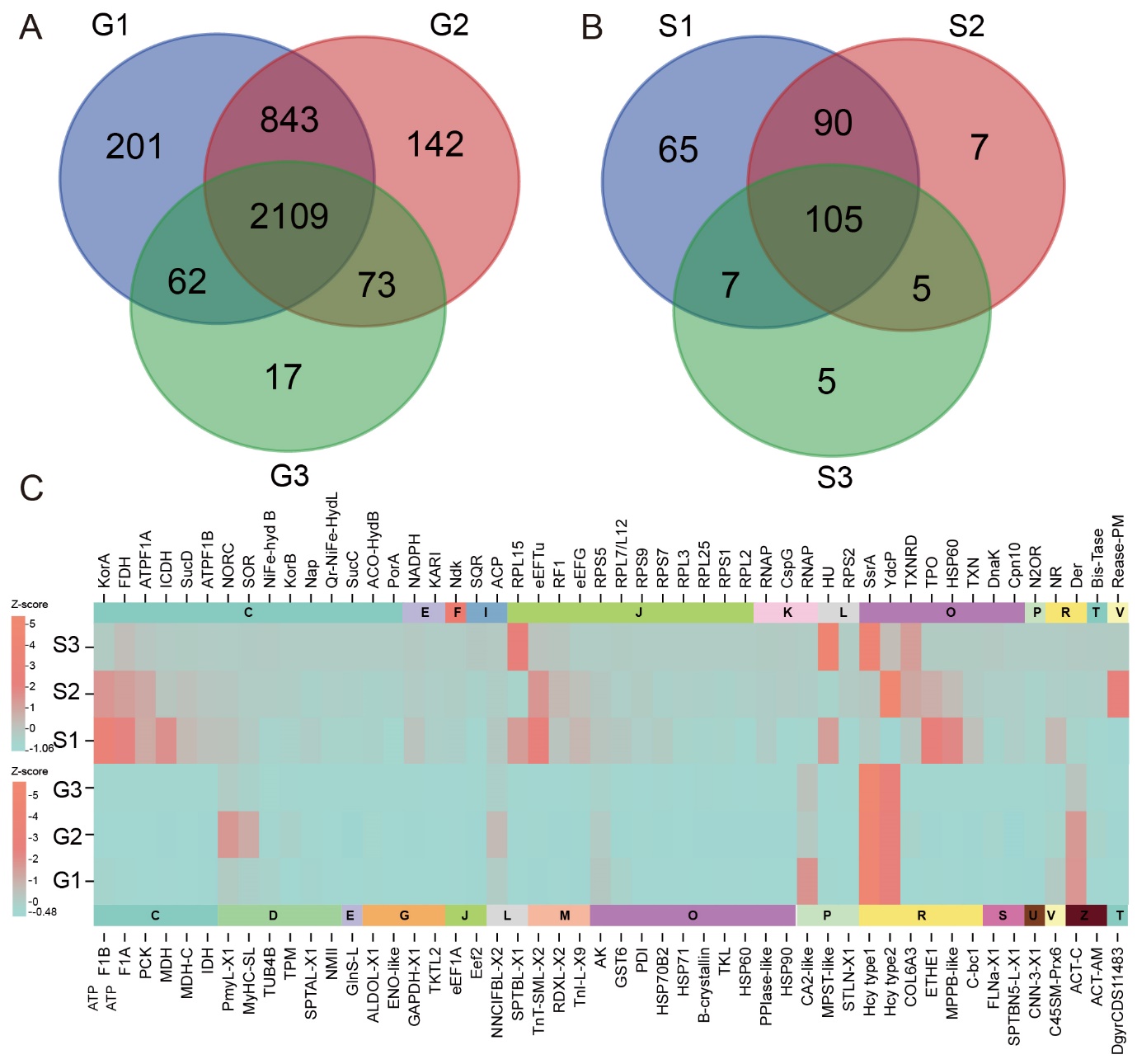

**Figure S6. *Alviniconcha marisindica* metaproteome analysis.** (A and B) Two venn diagrams of proteins identified of *Alviniconcha masindica* in three biological replicates. Host: G1, G2, G3; Symbiont: S1, S2, S3. (C) Heatmap of identified top 50 most abundant proteins in the metaproteomes. Abbreviations: S, symbiont; G, gill.

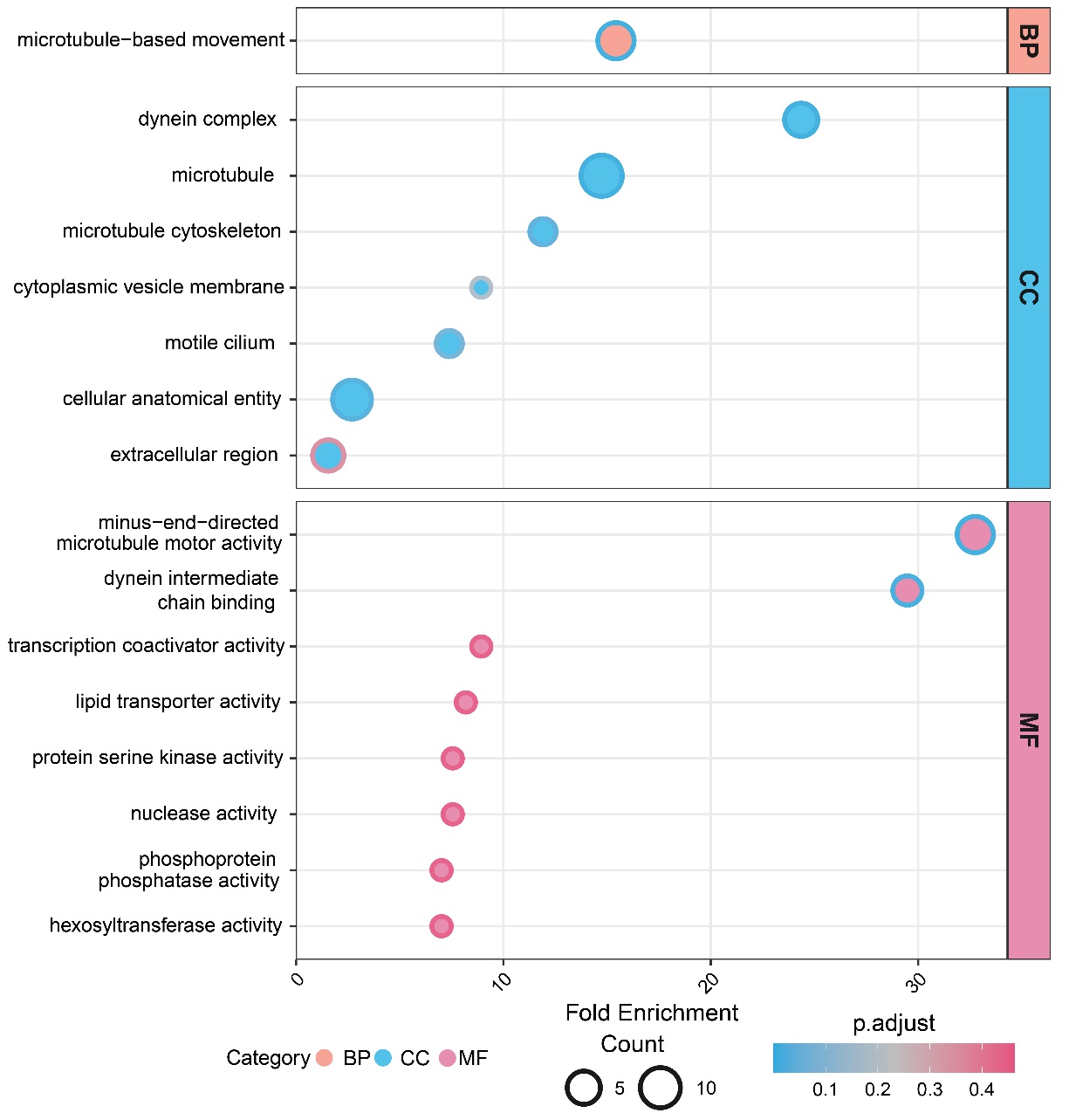

**Figure S7. GO enrichment of ciliated zone of *Alviniconcha marisindica* gill filaments as identified in the spatial transcriptomic analysis.**

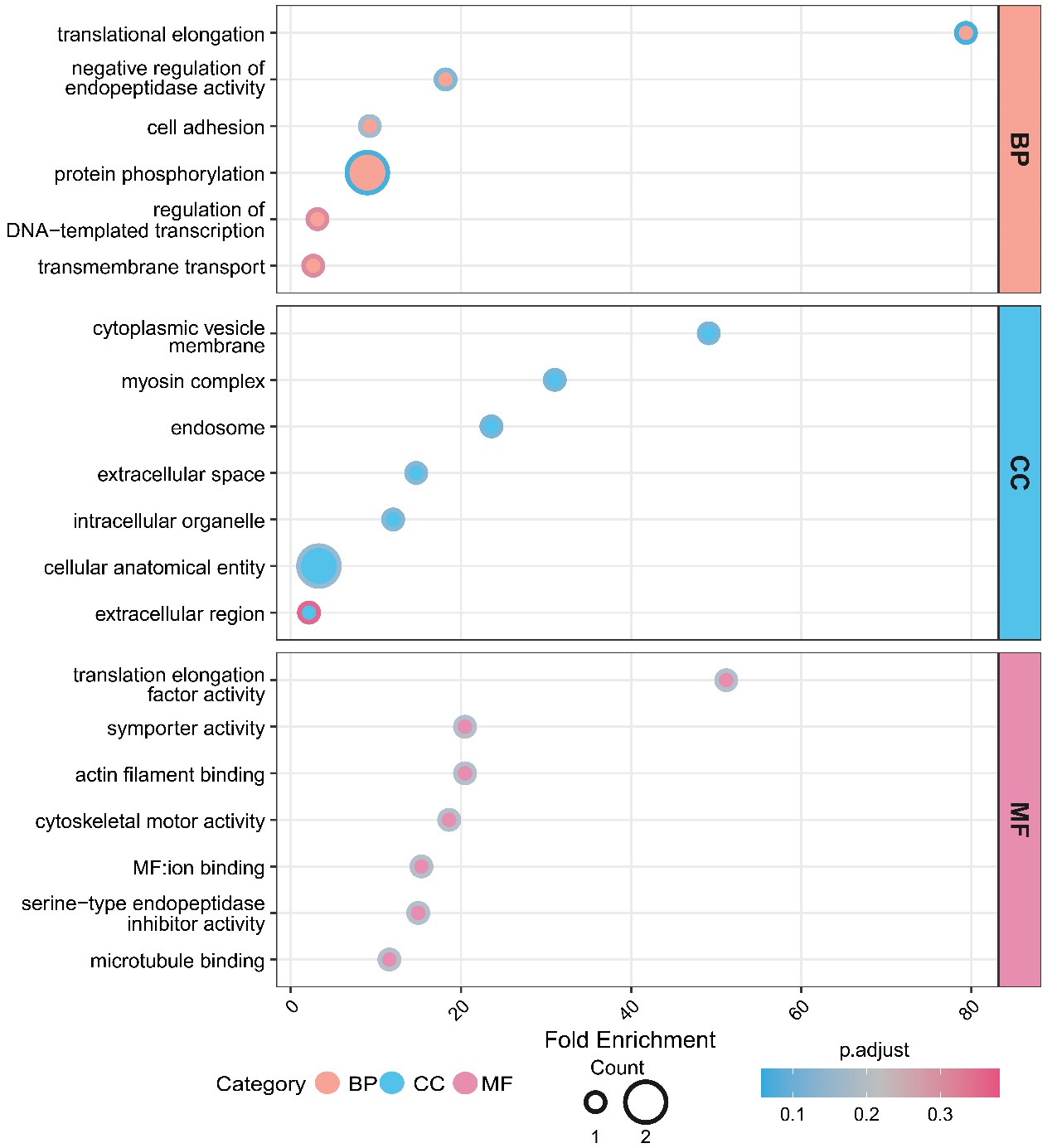

**Figure S8. GO enrichment of supporting axis of *Alviniconcha marisindica* gill filaments as identified in the spatial transcriptomic analysis**

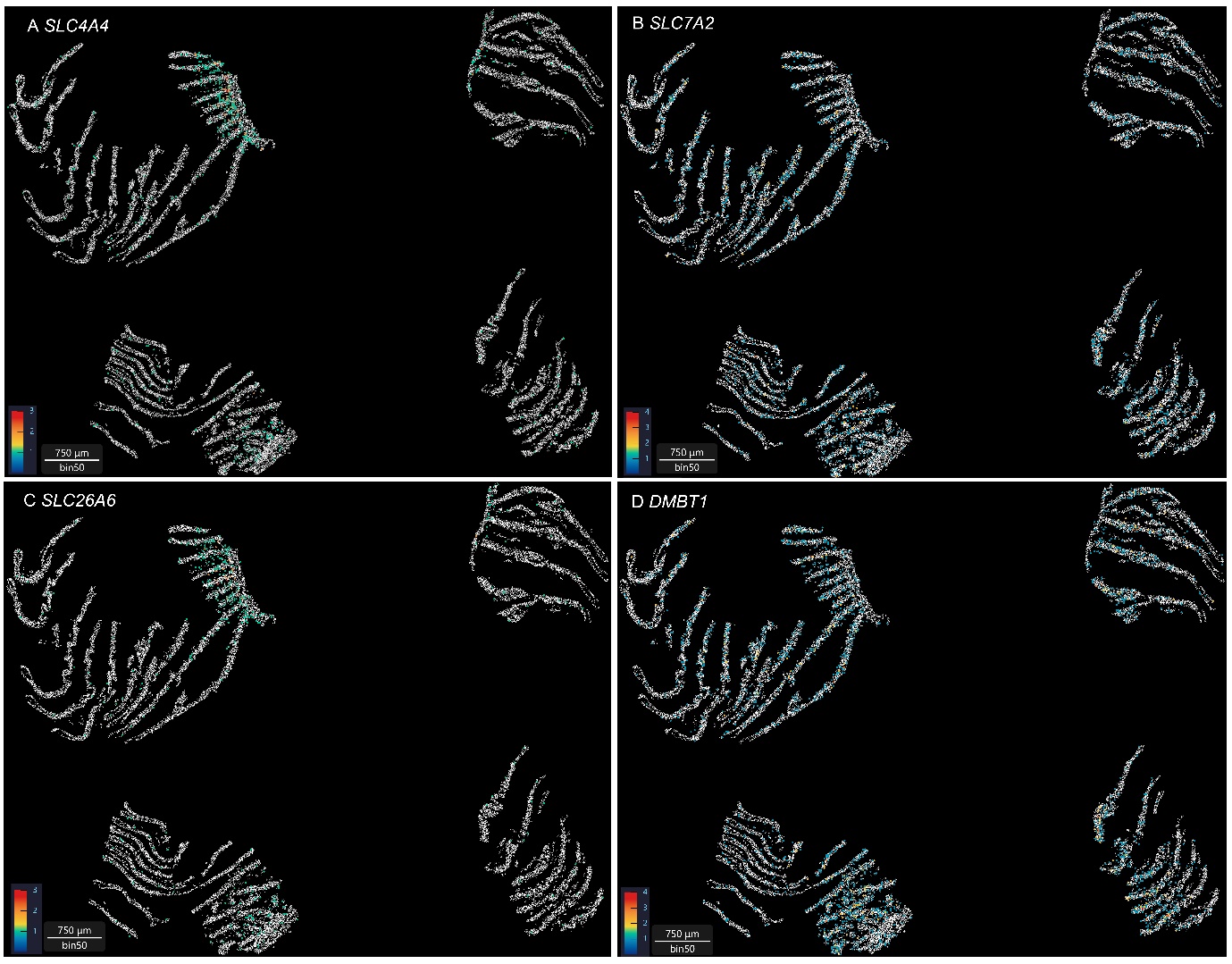

**Figure S9. Selected genes distribution pattern in *Alvinoncha marisindica* gill filaments** based on spatial transcriptome. (A) *SL4A4* distribution pattern in *A. marisindica* gill filaments. (B) *SLC7A2* distribution pattern in *A. marisindica* gill filaments. (C) *SLC26A6* distribution pattern in *A. marisindica* gill filaments (D) *DMBT1* distribution pattern in *A. marisindica* gill filaments.

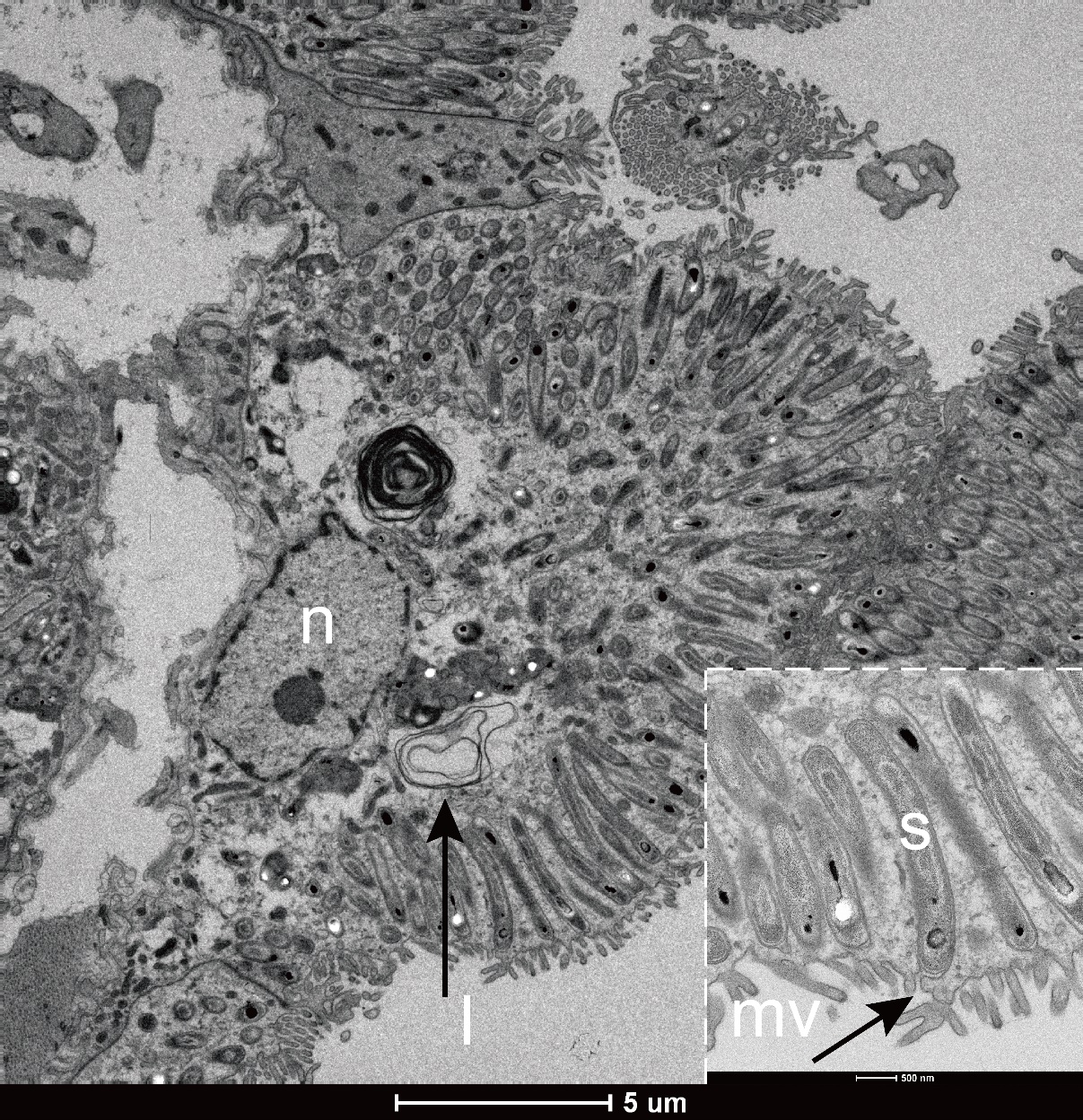

**Figure S10. Transmission electron microscopy analysis of gill section of *Alviniconcha adamantis*.** n: host nucleus; s: symbiont; l: lysosome; mv, microvilli. Scale bar: 5μm; 500nm.

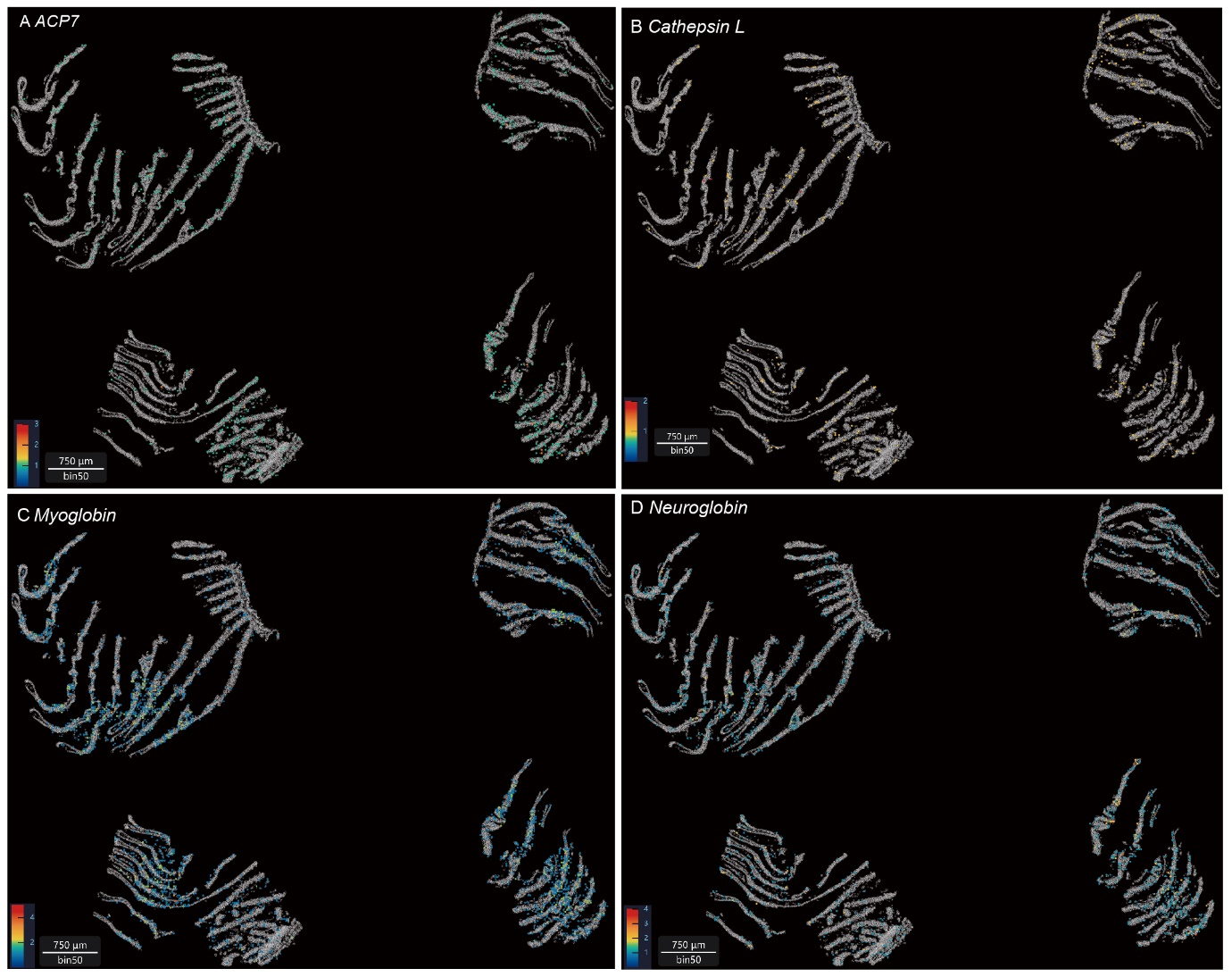

**Figure S11. Selected genes distribution pattern in *Alvinoncha marisindica* gill filaments** based on spatial transcriptome. (A) ACP7, acid phosphatase type 7, distribution pattern in *A. marisindica* gill filaments. (B) Cathepsin L distribution pattern in *A. marisindica* gill filaments. (C) Myoglobin distribution pattern in *A. marisindica* gill filaments (D) Neuroglobin distribution pattern in *A. marisindica* gill filaments.

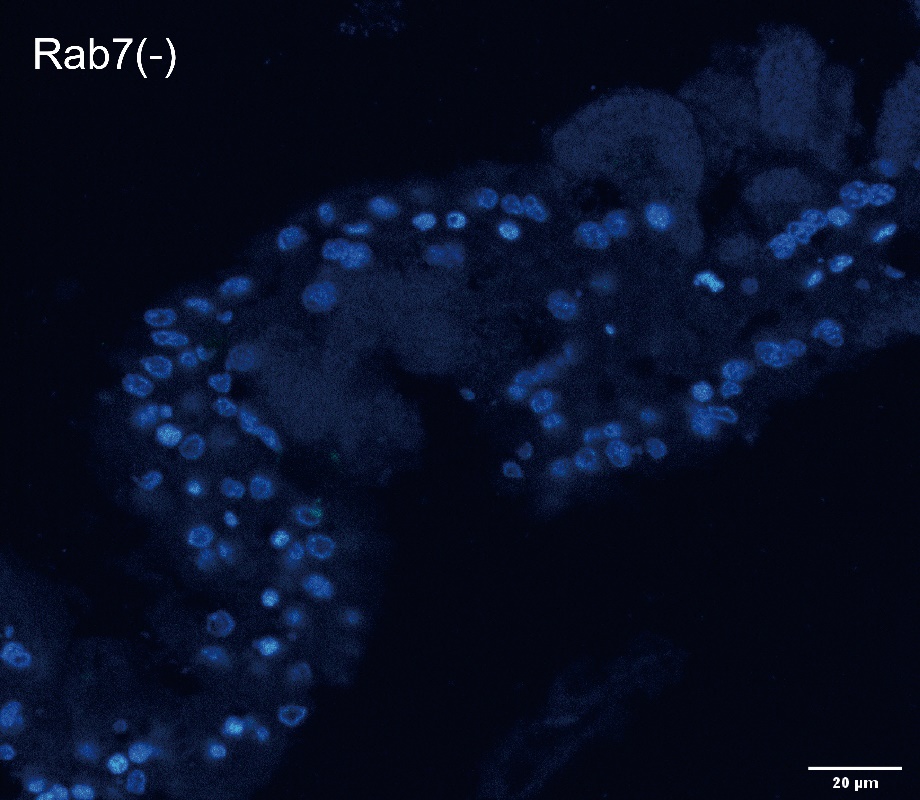

**Figure S12. Negative control of immunohistochemistry experiment using confocal microscope to detect Rab7 proteins in bacteriocyte zone of *Alviniconcha marisindica* gill filaments.** Blue fluorescent: nucleus. Scale Bar: 20μm.

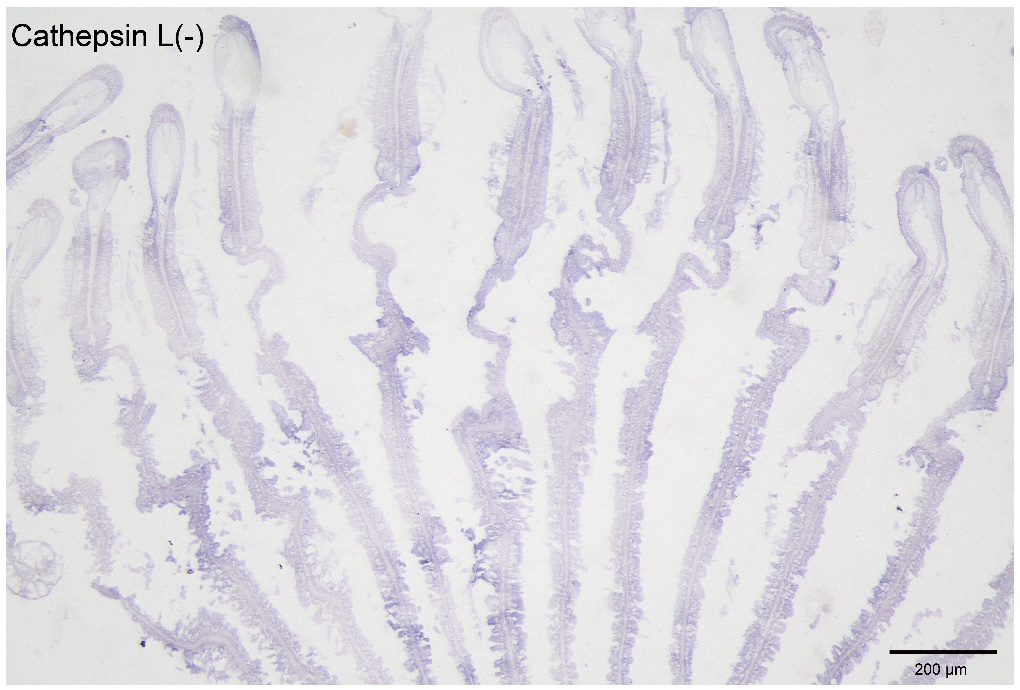

**Figure S13. Negative control of *in situ* hybridization to detect cathepsin L in the *Alviniconcha marisindica* gill filaments.** Scale bar: 200μm.

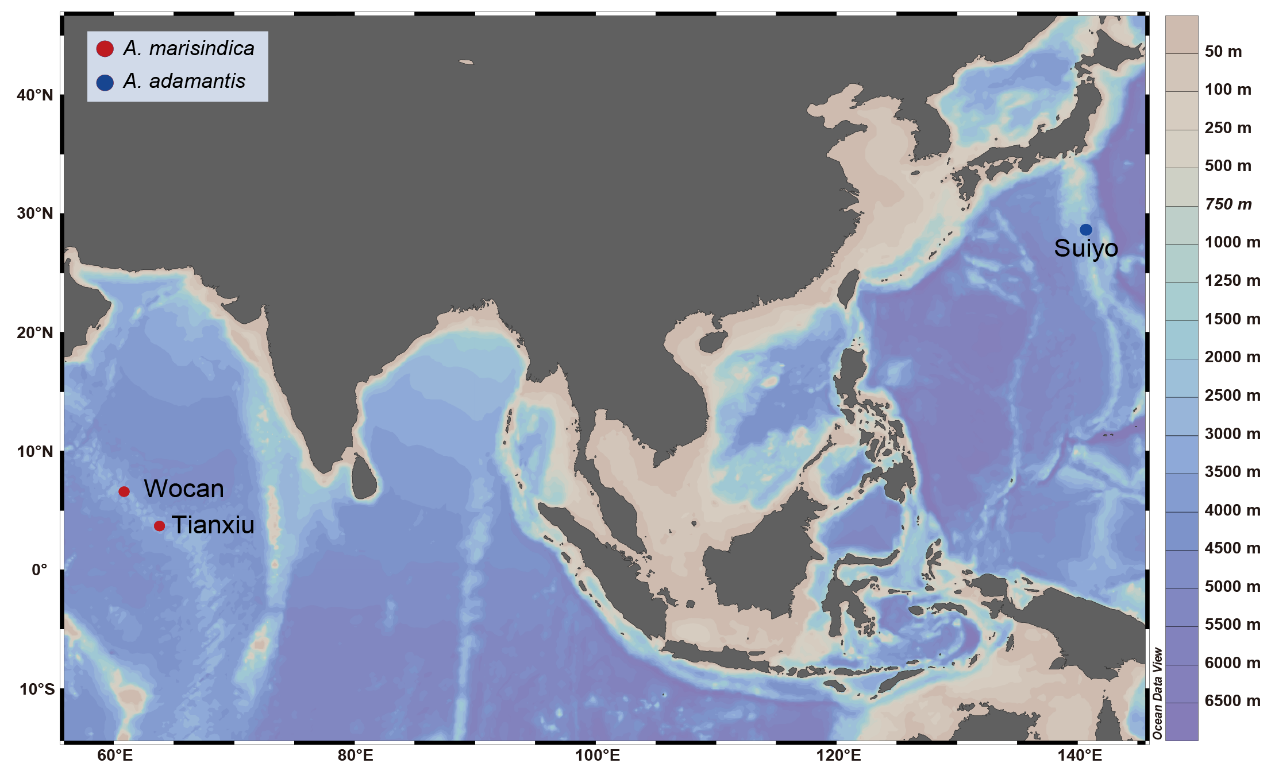

**Figure S14. Sampling localities of *Alvinoconcha marisindica* (Wocan and Tianxiu) and *Alviniconcha adamantis* (Suiyo) used in this study.**

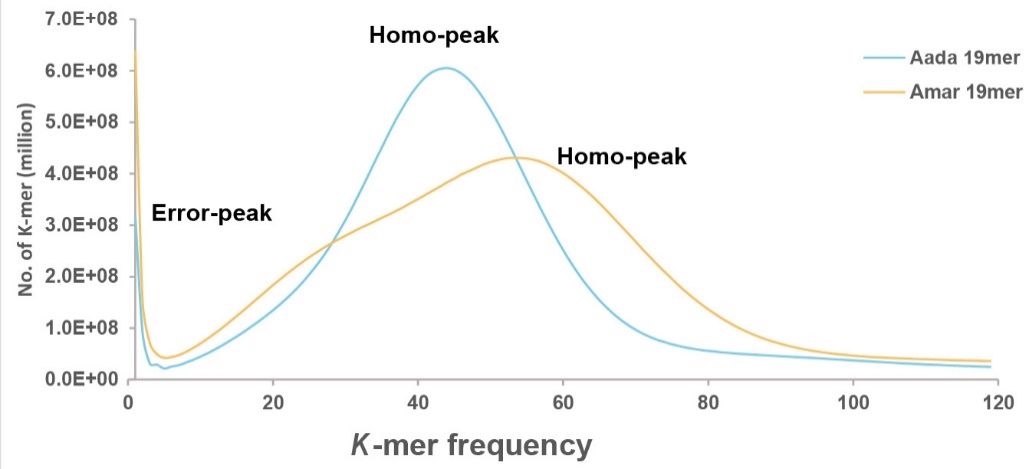

**Figure S15.** **Distribution of 19-mer frequency in two *Alviniconcha* species genomes.** High-quality sequencing reads generated from 150-bp libraries were used to generate the 19-mer depth distribution curve. Only one single homo-peak was identified in two species, indicating low genomic heterozygosity.

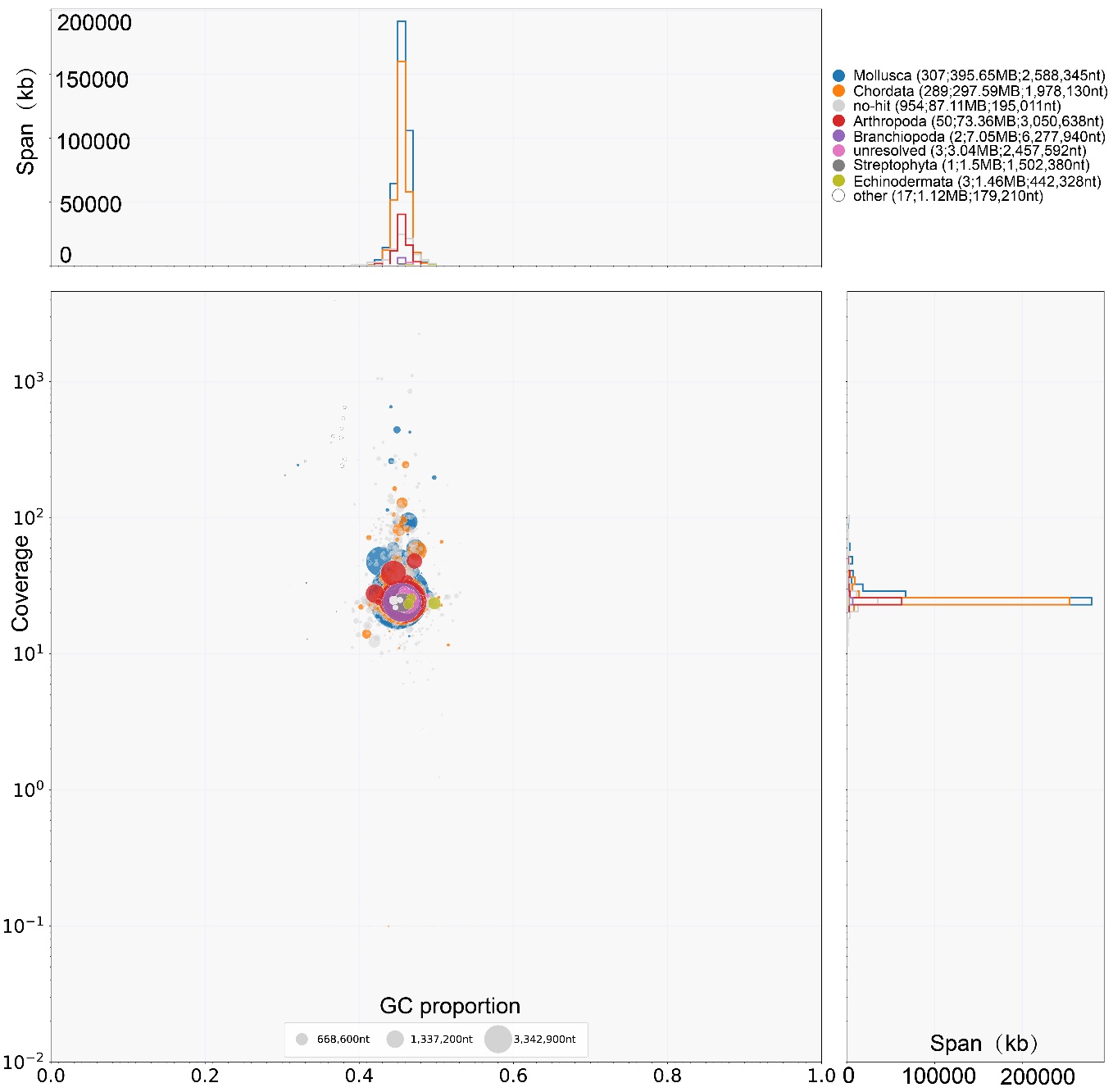

**Figure S16. Dot plot of *Alviniconcha* *marisindica* genome generated by Blobtools.** Since the neck tissue is non-symbiont tissue and was used for the genome sequencing, there are no microbe-contaminated reads in *A.* *marisindica.*

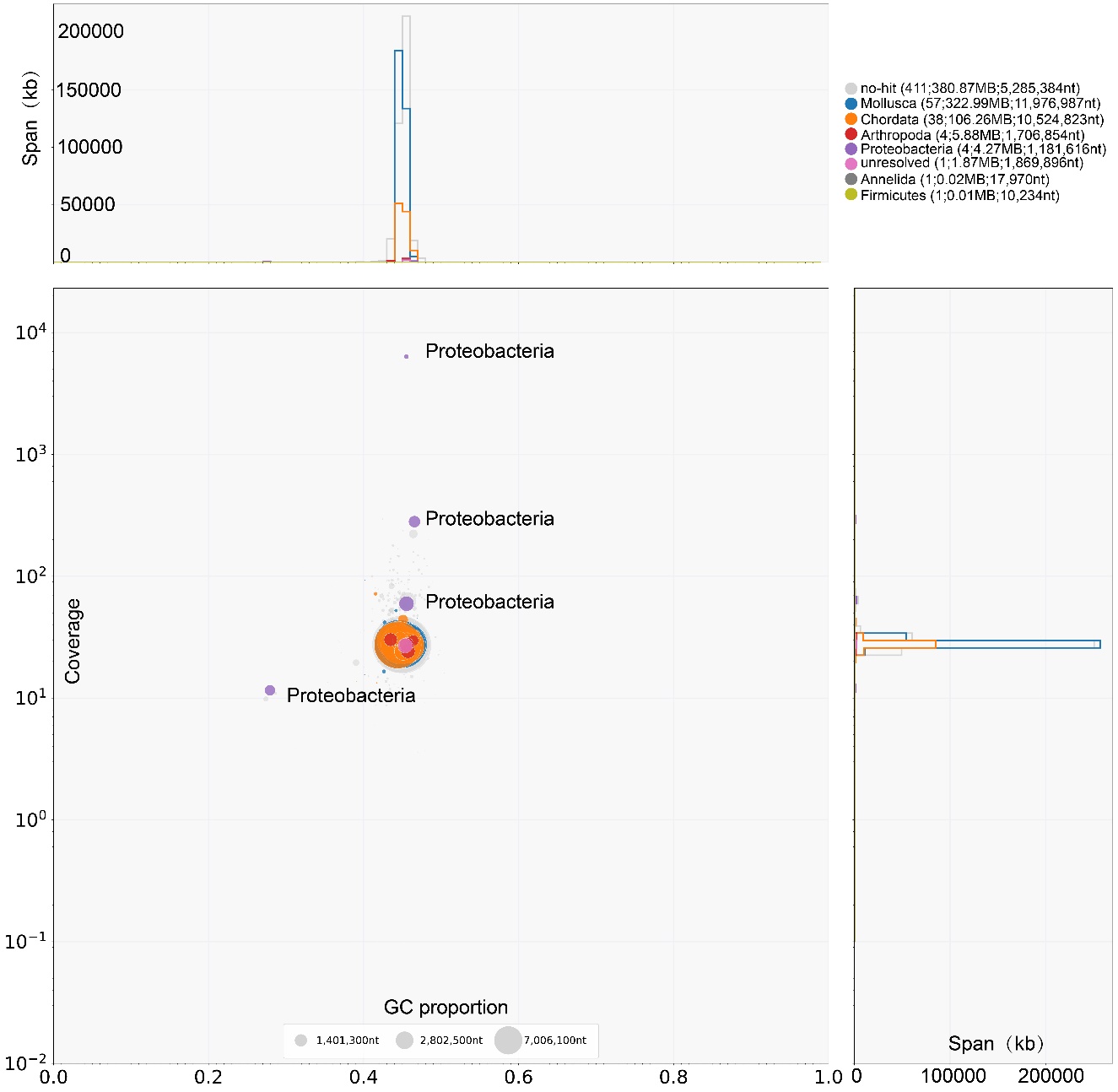

**Figure S17. Dot plot of *Alviniocncha adamantis* genome generated by Blobtools.** The purple circles are the microbe-contaminated contigs in *A. adamantis.*
